# Silencing the signal: The metastasis suppressor NDRG1 disrupts exosome-mediated crosstalk in pancreatic cancer

**DOI:** 10.1101/2025.10.15.682724

**Authors:** Jiawei Chang, Shafi Alenizi, Heloisa Zaccaron Milioli, Winston Lay, Saranya Pounraj, Yujie Li, Mekonnen Sisay Shiferaw, Elham Hosseini Beheshti, Zaklina Kovacevic

**Affiliations:** Department of Physiology, School of Biomedical Sciences, Faculty of Medicine and Health, University of NSW, Sydney NSW Australia 2052; School of Medical Sciences, Faculty of Medicine and Health, The University of Sydney, NSW 2006, Australia; Garvan Institute of Medical Research, Darlinghurst, NSW Australia 2010; Asbestos and Dust Disease Research Institute (ADDRI), Sydney, NSW, Australia 2138

**Keywords:** Pancreatic cancer, exosomes, NDRG1, ESCRT pathway, Alix, tumour microenvironment

## Abstract

Pancreatic cancer (PaC) remains one of the deadliest cancers, with 5-year survival rates of 13%. A major driver of its aggressiveness is the tumour microenvironment (TME), which fuels tumour growth, metastasis, and therapeutic resistance through dynamic, bi-directional communication between cancer cells, fibroblasts, and immune cells. Emerging evidence highlights extracellular vesicles (EVs) as key mediators of oncogenic cross-talk within the PaC TME. This study demonstrates for the first time that the metastasis suppressor NDRG1 significantly influences the biogenesis, cargo packaging and release of EVs by cancer cells. This was mediated by a direct interaction between NDRG1 and ALIX, a key protein involved in EV biogenesis and packaging, with NDRG1 facilitating ALIX proteasomal degradation. Further, EVs released from NDRG1-overexpressing cells had significantly fewer CAF-activation proteins (*i.e.* TGF-B), leading to attenuated ERKl/2 and p38 activation in pancreatic stellate cells (PSCs), and reduced expression of key fibrotic markers (a-SMA, FAP, and collagen 1A). NDRG1 also reduced EV uptake by PaC cells and diverted these to the lysosome for degradation. These findings uncover a previously unrecognized mechanism by which NDRG1 disrupts the oncogenic two-way communication between PaC cells and the TME, positioning NDRG1 as a compelling therapeutic target against this formidable malignancy.

**Graphical Abstract:** 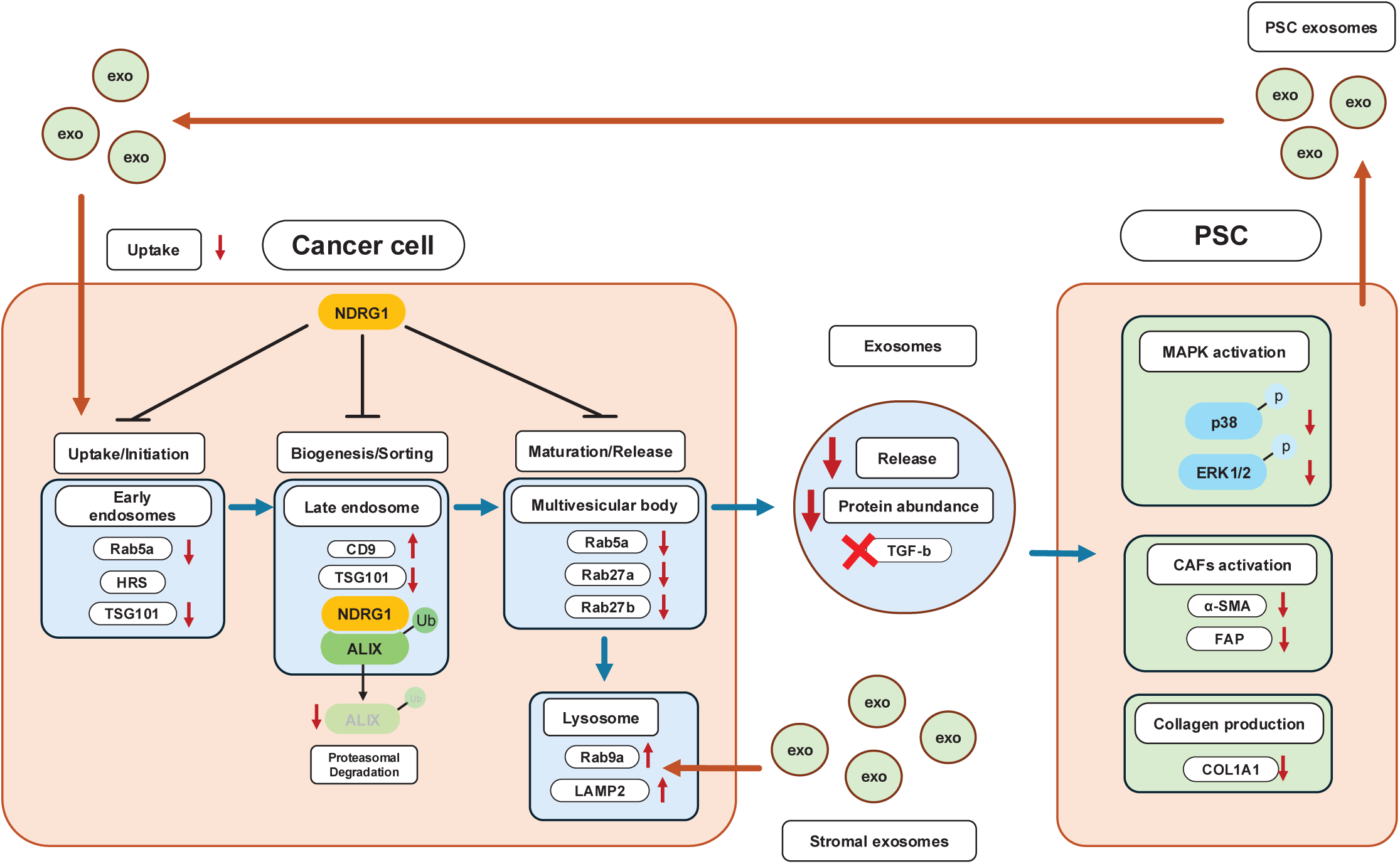

## 1. Introduction

Pancreatic cancer (PaC) is a highly aggressive malignancy associated with very poor 5-year survival rates of 13%. Current PaC treatments fail to significantly extend survival due to limited drug penetration into the hypovascularized and dense PaC tumour microenvironment (TME) [1]. Over the past decade, extracellular vesicles (EVs) have emerged as critical mediators of intercellular communication within the cancer TME. These nanosized membrane-encapsulated vesicles contain bioactive materials such as nucleic acids, fatty acids, proteins, and metabolites that are secreted by almost all cell types [2]. Cancer cells release EVs through different biogenesis pathways to interact with cells within the TME and extracellular matrix (ECM), creating a favourable niche for proliferation and invasion [3]. EVs interact with other cells in local and distant locations, potentially altering the activity of recipient cells [4]. In PaC, EVs were found to facilitate cancer cell migration and invasion [5] and to prepare the pre-metastatic niche at distal sites such as the liver [6]. Mounting evidence suggests that the oncogenic progression of PaC is tightly associated with cellular alterations caused by EVs [3, 7].

Bidirectional communication between cancer and stromal cells *via* EVs drives multiple aspects of PaC progression. Pancreatic stellate cells (PSCs), which are abundant in the pancreas and a key pre-cursor to cancer-associated fibroblasts (CAFs;[8]) release EVs containing miRNA cargo that promote PaC cell proliferation, migration, and metastasis in both cell and animal models [9–13], while also contributing to gemcitabine resistance [14, 15]. Conversely, PaC cell-derived EVs alter the function of neighbouring CAF and immune cells in the TME to promote desmoplasia and immune escape [16, 17]. These EVs contain bioactive materials that can initiate the activation of PSCs into CAFs, promote CAF collagen production, and polarize local macrophages into the oncogenic M2 phenotype [3, 15].

Exosomes, the most extensively studied subtype of EVs, undergo a tightly regulated biogenesis process involving two main regulatory pathways: the endosomal sorting complex required for transport (ESCRT) pathway and small GTPase Rab family protein-facilitated vesicle trafficking [l8-20]. Manipulation of these pathways in cancer cells can significantly impair exosome release and cargo sorting capacity [21–23]. It has been observed that the absence of Annexin A1 (ANXA1), a key promoter and regulator of PaC development and drug resistance [24–26], dramatically decreases the quantity of secreted exosomes by PaC cells [10]. Among the key components of the ESCRT machinery, ALG-2-interacting protein X (ALIX) and tumor susceptibility gene 101 (TSG101) play major roles in exosome biogenesis. ALIX functions as an ESCRT-associated protein that facilitates membrane budding and cargo selection [27], while TSG101 is a core component of ESCRT-I that mediates the sorting of ubiquitinated proteins into intraluminal vesicles [28]. Notably, ALIX and TSG101 have been shown to interact directly or within a shared complex, coordinating the formation of intraluminal vesicles (ILV) within multivesicular bodies (MVBs) [29]. This interaction underscores their cooperative function in regulating ESCRT-dependent exosomal pathways and supports their classification as hallmark markers and effectors of exosome production. Overall, inhibition of EV biogenesis can potentially disrupt communication between PaC and its TME, thus hindering tumour progression and metastasis.

Recent studies have identified that the metastasis suppressor N-myc downstream-regulated gene 1 (NDRG1) can potently inhibit the cross-talk between PaC cells and stromal PSCs [30, 31]. Intriguingly, NDRG1 has also been suggested to influence endosomal sorting and EV biogenesis [32–34], although its effects on PaC EV secretion and EV cargo packaging have never been assessed. NDRG1 is commonly expressed in the cytoplasm and nuclei of epithelial tissues and was identified as a metastasis suppressor [35]. In normal cells, NDRG1 regulates multiple cellular pathways and processes relating to differentiation [36, 37], lipid synthesis [38], and apoptosis [39]. Accumulating evidence suggests that NDRG1 plays an important inhibitory role in the progression and metastasis of PaC [40–43]. This is underscored by studies showing how NDRG1 inhibits tumour growth [44], migration and metastasis [45], angiogenesis [46] and a plethora of oncogenic signalling pathways in PaC cells [30, 31, 47, 48].

Recent studies have reported that NDRG1 expression in cancer cells can also influence the TME [49, 50]. In PaC cell models, Geleta *et al.* demonstrated that upregulation of NDRG1 can suppress the formation of desmoplasia by disrupting the communication between PSCs and PaC cells, with novel anti-cancer agents that up-regulate NDRG1 effectively inhibiting PaC growth and metastasis *in vivo* [30, 31]. However, the mechanisms by which NDRG1 inhibits PaC-PSC communication remain to be elucidated.

In the current study, we demonstrate for the first time that NDRG1 functions as a master regulator of exosome biogenesis in PaC. Our findings reveal that NDRG1 significantly reduces exosome secretion by directly interacting with and promoting the proteasomal degradation of ALIX, a key component of the ESCRT machinery. This interaction substantially alters exosomal cargo composition, particularly depleting tumour-promoting factors such as TGF-Bl. Furthermore, NDRGl overexpression attenuates PSC activation by exosomes, reduces CAF marker expression, and limits desmoplastic reactions. Additionally, we show that NDRG1 also reduces cancer cell uptake of TME-derived exosomes while promoting their lysosomal degradation. Together, these findings establish a novel mechanism by which NDRG1 inhibits tumour-stroma cross-talk in PaC and reveals NDRG1 as a potential therapeutic target to modulate exosome-mediated influence on the TME.

## 2. Materials and methods

### 2.1 Cell culture

The human pancreatic cancer cell lines PANC-1 and MIAPaCa-2 were purchased from the American Type Culture Collection (Rockville, MD). Cells were authenticated based on viability, recovery, growth, morphology, and cytogenetic analysis by the provider. PANC-1 and MIAPaCa-2 cells were both derived from the epithelial cells of pancreatic carcinomas. Pancreatic stellate cells (CAT#: 3830) were purchased from ScienCell Research Laboratories (Carlsbad, CA). PANC-1 and MIAPaCa-2 cells were grown in Dulbecco’s modified Eagle’s medium (DMEM) (Gibco, CAT#:11965092). They were supplemented with l0% (v/v) foetal calf serum (FCS), l% (v/v) nonessential amino acids, l% (v/v) sodium pyruvate, l% penicillin/streptomycin (Invitrogen). In addition, MIAPaCa-2 cells were also supplemented with 2.5% horse serum (Gibco). PSCs were cultured with Iscove’s Modified Dulbecco’s Medium (IMDM), supplemented with l0% FCS and l% penicillin/streptomycin. Culture media were tested for mycoplasma every 3 months. All cells were grown in a tissue culture incubator at 37°C with 5% CO_2_ and sub-cultured by standard methods.

### 2.2 Transfection

Both PANC-1 and MIAPaCa-2 cells were transfected with pcDNA3.1MycHisA(-) plasmids containing wild type NDRG1 (NDRG1), which were kindly provided by Prof. Yoel Sadovsky (University of Pittsburgh, Pennsylvania; [51]). Empty pcDNA3.1MycHisA(-) plasmids were used as vector controls (VC; Invitrogen). To silence NDRG1, MIAPaCa-2 cells were transfected with an NDRG1 shRNA Plasmid (NDRG1*^KD^*) or shRNA negative control plasmid (NC; Abbexa, USA). All stably transfected cells were maintained in the presence of G4l8 (300 µg/ml) as described previously [25, 48].

### 2.3 EV isolation

To isolate EVs, PaC or PSC cells were first grown in their normal complete media for 48 h until 75% confluency, washed with warm PBS, followed by 48 h culture in FCS-free media, after which the overlaying media was collected. This condition media (CM) then underwent 3 x 300 RCF (5 min) centrifugation steps at 4°C to remove cell debris, followed by l x 2800 RCF (10 min) centrifugation at 4°C to isolate large oncosomes and apoptotic bodies. The supernatant then underwent gradient ultracentrifugation at 10,000 RCF at 4°C for 45 min to isolate 10K EVs (Large EVs). Exosomes were then collected by ultracentrifuging the remaining supernatant at 100,000 RCF at 4°C for 65 min (**Figure 1A**). EV pellets were suspended in 200 µL of exosome-free PBS or 100uL of RIPA buffer (for protein extraction) and stored at −80°C until further use. These methods for enriching different EV populations are well established [52–56]. Protein was extracted from remaining cancer cells using protein lysis buffer for Western blotting or Pierce™ IP Lysis Buffer (Thermo Scientific) for co-immunoprecipitation.

**Figure 1.**
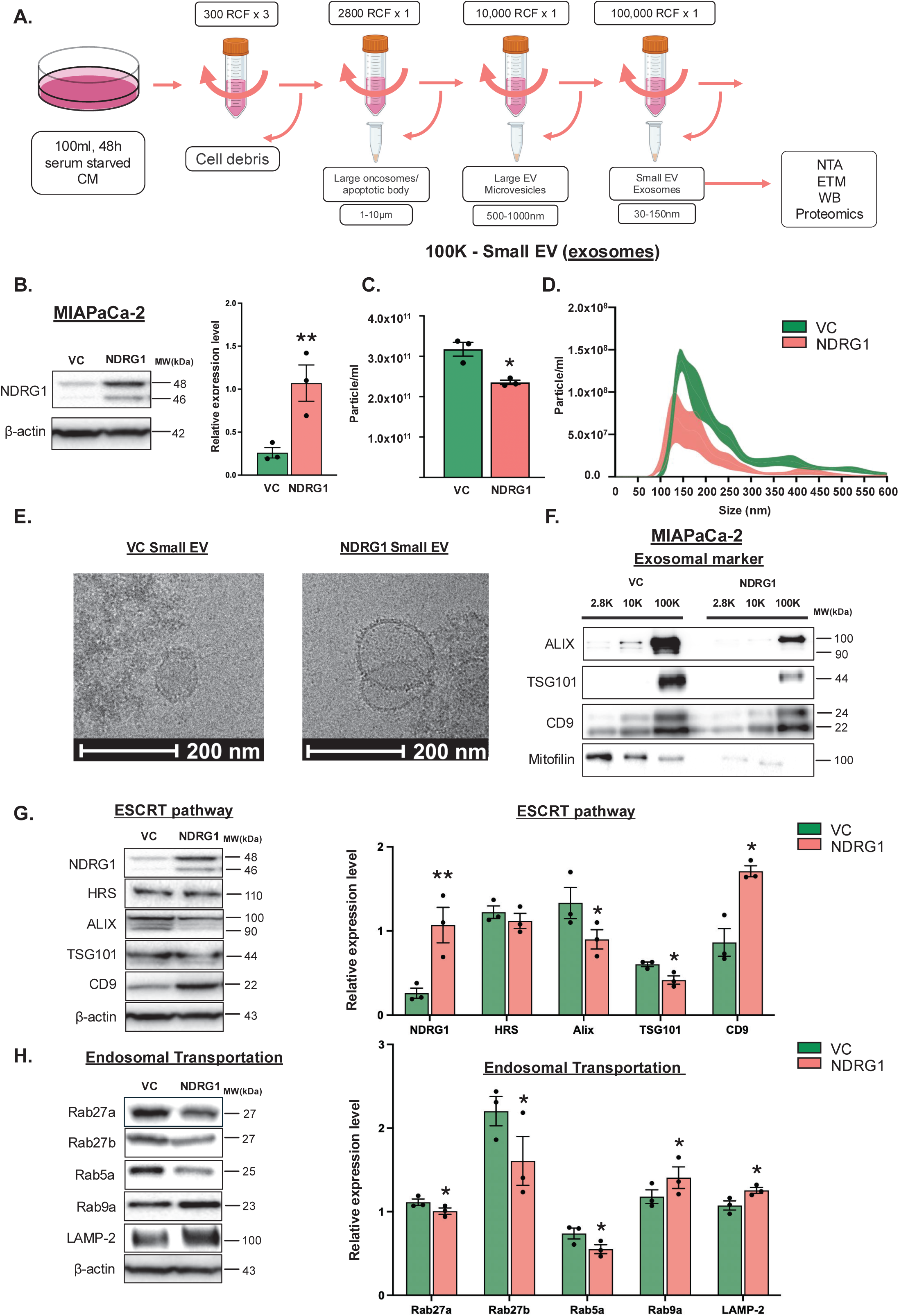
NDRG1 reduces extracellular vesicle (EV) production and biogenesis pathways in pancreatic cancer cells. (**A**) Schematic of the gradient ultracentrifugation procedure used to enrich large and small extracellular vesicles (EV) from the conditioned medium (CM) of serum-starved MIAPaCa-2 cells. (**B**) Representative western blot images showing NDRG1 expression in MIAPaCa-2 vector control (VC) cells and MIAPaCa-2 cells stably transfected to over-express NDRG1 (NDRG1), with B-actin as a loading control. The accompanying densitometry analysis quantifies relative NDRG1 expression, with each dot representing an independent experiment (n=3). (**C**) Nanoparticle tracking analysis (NTA) showing the concentration of small EVs (exosomes) isolated from VC or NDRG1 cells with each dot representing an independent experiment. (**D**) The size distribution graph depicts the particle size profile of exosomes measured by NTA, with shaded regions representing the standard error of the mean (SEM), calculated from three independent experiments. (**E**) Representative transmission electron microscopy (TEM) images of small EVs isolated from MIAPaCa-2 VC and NDRG1 cells. Scale bar: 200 nm. (**F**) Representative western blot images of exosomal markers (ALIX, TSG101, and CD9) and mitochondrial marker (Mitofilin) in 2.8K, 10K, and 100K EV fractions isolated from MIAPaCa-2 VC and NDRG1 cells. Representative western blot images of (**G**) ESCRT pathway proteins (HRS, ALIX, TSG101, and CD9) and (**H**) endosomal trafficking regulators (Rab27a, Rab27b, Rab5a, Rab9a, and LAMP-2) in VC or NDRG1 MIAPaCa-2 cells. The bar graphs adjacent to the blots represent the densitometric analysis, which was normalized to B-actin. Each dot represents an independent experiment, and bars indicate the mean ± SEM. Statistical significance was determined using Student’s *t*-test and is indicated as \**p* < 0.05; \*\**p* < 0.01.

### 2.4 NanoSight^TM^ particle tracking analysis and ZetaSizer ultra sizing

Isolated exosomes were analysed using the NanoSight NS300 nanoparticle characterisation system (Nanosight Ltd., UK) for their size and concentration. EV samples were diluted in particle free PBS to a measurable concentration between 0.5×10^8^ and 5×10^9^ particles/ml. During the analysis, 1 ml of EV samples were injected into a viewing unit using a controlled syringe system. In the viewing unit, a monochromatic laser beam (488 mm) was applied to the diluted samples. 5 independent recordings were taken per sample for concentration and size distribution analysis. MIAPaCA-2 VC and NDRG1 derived EVs with same dilution factors were also examined with Malvern ZetaSizer Ultra (ATA Scientific Pty Ltd, AU) for sizing of EVs.

### 2.5 Exosome Cryo-EM data acquisition

EV samples (4.5 µL) were applied to glow discharged Quantifoil R2/2 copper grids (Quantifoil Micro Tools) and blotted for 2.5 s in a 95% humidity chamber, then plunged in liquid ethane using a Leica EM GP device (Leica Microsystem). The grids were imaged using a Talos Arctica cryoTEM (Thermo Fisher Scientific) and operated at 200kV, with the specimen maintained at liquid nitrogen temperatures. Images were recorded on a Falcon 3EC direct detector camera operated in linear mode.

### 2.6 Protein extraction

PaC cells were cultured in FCS-free media for 48 h prior to protein extraction. Protein extraction was performed as described previously [57], using freshly-made lysis buffer (10 mM Tris buffer, 150 mM NaCl, 5% SDS, 1% Triton X-100, 1 mM EDTA, 0.04 mM NaF, 4% protease inhibitor (Roche, Germany) and 2x PhoSTOP (Roche, Germany)). Extracted lysates were sonicated on ice followed by centrifugation at 4°C, 14,000 RPM for 40 min. The supernatant was stored at −80°C. Protein concentration was measured using the BCA protein assay (Thermo Scientific). EV protein concentration was measured using a microBCA kit (Thermo Scientific). Protein absorbance was measured at 562 nm.

### 2.7 Western blotting

Western analysis was performed *via* standard procedures [48] using antibodies listed in **Table 1**. Western blots were imaged using Immobilon ECL Ultra Western HRP Substrate (MERCK, CAT#: WBULS0100) with the Bio-Rad ChemiDoc MP Imaging System (Bio-Rad. Australia). Band intensity was analysed with the Bio-Rad Image Lab software.

**Table 1:** Antibodies used for immunoblotting and immunofluorescence. Primary and secondary antibodies used for western blotting (WB) and immunofluorescence (IF), including the working dilutions for each application. All antibodies were used according to the manufacturer’s instructions, with dilutions optimized for analysis.

| **Antibody** | **Catalogue #** | **Company** | **Dilution (WB)** | **Dilution (IF)** |
| --- | --- | --- | --- | --- |
| NDRG1 | 9485S | Cell signalling technology | 1:1000 | 1:200 |
| NDRG1 | J6271-2D7 | Sigma-Aldrich |  | 1:200 |
| ALIX | 92280 | Cell signalling technology | 1:1000 | 1:800 |
| Ubiquitin | E4I2J | Cell signalling technology | 1:1000 |  |
| RAB5A | 46449 | Cell signalling technology | 1:1000 | 1:200 |
| RAB27A | 69296 | Cell signalling technology | 1:1000 | 1:200 |
| RAB27B | 17572 | Cell signalling technology | 1:1000 | 1:500 |
| RAB9A | 95394 | Cell signalling technology | 1:1000 | 1:200 |
| HRS | A300-989A-M | Bethyl Laboratories | 1:1000 |  |
| TSG101 | ab30871 | Abcam | 1:1000 |  |
| CD9 | 13403 | Cell signalling technology | 1:1000 |  |
| LAMP2 | ab25631 | Abcam | 1:500 | 1:500 |
| TGF-β1 | ab215715 | Abcam | 1:1000 | |
| aSMA | 19245 | Cell signalling technology | 1:1000 |  |
| FAP | 66562 | Cell signalling technology | 1:1000 |  |
| COL1A1 | 72026 | Cell signalling technology | 1:1000 | 1:1000 |
| p-ERK1/2 (Thr202/Tyr204) | 9101 | Cell signalling technology | 1:500 |  |
| p-P38a (Thr180/Tyr182) | 9211 | Cell signalling technology | 1:500 |  |
| Total ERK1/2 | 4695 | Cell signalling technology | 1:1000 |  |
| Total P38a | 8690 | Cell signalling technology | 1:1000 |  |
| $\beta$ - actin | BA3R | Thermo Scientific | 1:10000 | |
| Anti-rabbit IgG, HRP-linked Antibody | 7074 | Cell signalling technology | 1:10000 |  |
| Goat anti-Mouse IgG (H+L) Secondary | 31430 | Thermo Scientific | 1:10000 |  |
| Anti-rabbit IgG (H+L), F(ab')<sub>2</sub> Fragment (Alexa Fluor® 594 Conjugate) | 8889 | Cell signalling technology |  | 1:500 |
| Anti-mouse IgG (H+L), F(ab')<sub>2</sub> Fragment (Alexa Fluor® 488 Conjugate) | 4408 | Cell signalling technology |  | 1:500 |

### 2.8 Immunofluorescent staining and microscopy

Transfected MIAPaCa-2 cells were seeded into 24-well plates onto sterile 1 cm round coverslips. Cells were incubated in their standard media for 24 h, followed by a further 48 h incubation in FCS-free media. Cells were fixed with 4% paraformaldehyde (PFA) in PBS for 10 min, washed and permeabilized with 0.1% Triton X-l00/PBS for l5 min at room temperature. After blocking with 5% BSA/PBS (45 mins), the cells were incubated with primary antibody diluted in l% BSA/PBS overnight at 4°C. Antibodies used and their concentration can be found in **Table 1**. After washing, cells were incubated with secondary antibodies at a dilution of l:500 in l% BSA/PBS for l h, washed with 0.2% BSA/PBS (3 x l0 min), stained with DAPI for 3 mins (NucBlue Fixed Cell ReadyProbes Reagent (DAPI), CAT#: R37606, Invitrogen) and mounted onto glass slides using mounting medium (ProLong™ Diamond Antifade Mountant, CAT#P36961, Invitrogen) and allowed to cure at room temperature for 24 h prior to imaging. Images were acquired with the Leica stellaris 8 confocal imaging system.

### 2.9 Proximity ligation assay

VC and NDRG1 overexpressing MIAPaCa-2 cells were seeded on to 8-well chamber slides (CAT#: 154534, ThermoFisher Scientific). Once cells reached 75% confluency, they were cultured in FBS free media for 48 h. Cells were fixed with 4% PFA at room temperature for 10 min, then they were washed and permeabilized with 0.1% Trion-X-l00/PBS for l5 min. Proximity ligation assay was performed using the Duolink® In Situ Red Starter Kit Mouse/Rabbit (CAT#: DUO92101, Sigma-Aldrich) following the manufacturer’s protocol using ALIX (CAT#: 92280, Cell Signalling technology, l:800 dilution) and NDRG1 (CAT#: J6271-2D7, Sigma-Aldrich, 1:200 dilution) antibodies. Images were taken on Leica stellaris 8 confocal imaging system with 20x magnification.

### 2.10 Co-immunoprecipitation and MG132 protein degradation assessment

Protein lysates were extracted from MIAPaCa-2 VC and NDRG1-overexpressing cells using Pierce™ IP Lysis Buffer supplemented with 4% protease inhibitor cocktail (Roche, Germany), after serum starvation for 48 h. For immunoprecipitation, NDRG1 Rabbit monoclonal antibody (CAT #: 9485S, Cell Signalling Technology, 1:50 dilution), ALIX Rabbit monoclonal antibody (CAT #: 92880S , Cell Signalling Technology, 1:50 dilution), or Rabbit IgG isotype control (Cell Signalling Technology, CAT#: 3900, concentration matched to antibody) were incubated with Dynabeads™ (Thermo Scientific) for 4 h at 4°C on a rotating mixer. Subsequently, 30 μL of bead-antibody complexes were incubated with 200 µL of protein lysate (2 µg/µL) overnight at 4°C with rotation. Prior to Western blotting, samples were thoroughly washed with IP Lysis Buffer. Input controls were taken before bead-antibody incubation, and supernatant controls (l5 µL, 2 µg/µL) were collected post-incubation. To assess ALIX ubiquitination, cells were treated with 2.5 μM MG132 (CAT#: 2194, Cell Signalling Technology,) for 24 h prior to extraction and Co-IP. Western blotting analysis employed ALIX antibody (CAT#: 92880S, Cell Signalling Technology, diluted l:l000), NDRGl antibody (Abcam, Cat#: ab37897, diluted 1:1000), and ubiquitin antibody (CAT#: E4I2J, Cell Signalling Technology, diluted 1:1000), following the method described in *Section 2.7*.

### 2.11 Cycloheximide Chase Protein half-life assay

MIAPaCa-2 VC and NDRG1 cells were treated with cycloheximide (CHX, 15 µg/mL; CAT#: 2112, Cell Signalling) diluted in serum free media to inhibit protein synthesis. CHX treatment was administered for 0, 4, 8, l6, 24, 36, and 48 h, followed by protein extraction and western blotting.

### 2.12 NDRG1 and ALIX protein remodelling and docking

The 3D models of NDRG1, ALIX and their predicted interaction were generated using AlphaFOLD3 [58]. Amino sequence data were obtained from publicly available UniProt. The lowest energy state model was chosen for optimal representation. Docked profile and polar interaction were then visualised using PyMOL.

### 2.13 Label-free qualitative proteomic and DIA-NN library-free peptide search

Exosome and 10K EV samples derived from MIAPaCa-2 VC and NDRG1 cells were suspended in 1x RIPA buffer (Abcam) with 12.5% protease and phosphatase inhibitors (Roche, Germany). Quantitative proteomic analysis of cancer cell derived exosomes was performed by Sydney Mass Spectrometry using standard procedures [59]. Exosome samples were run on DIA mode on HFX4 for peptide search. The HFX4 output files were uploaded to DIANN version l.8.l and processed in a library free search against Human genome for peptide identification.

### 2.14 EV Proteomic Data Analysis

Raw proteomic data generated using DIANN was initially filtered to remove missing values, retaining only proteins identified in all three biological replicates of both MIAPaCa-2 VC and NDRG1-derived EVs. The filtered dataset was then log10-transformed to normalize protein abundance distributions. Differentially expressed proteins were identified using volcano plot analysis (*p*<0.01 and Fold change (FC) = 0 ) and heatmapping (*p* <0.05, ranked by FC), Finally, a t-SNE plot was generated in R Studio to visualize and cluster the samples based on their proteomic profiles.

### 2.15 Phospho-Kinase Array assay

PSCs seeded in 10 cm culture plates were incubated with MIAPaCa-2 VC and NDRG1-derived exosomes (2 x 10^10^ particles) for 1 h in serum free media, followed by protein extraction of the PSC lysate. The phosphorylation status of 37 human kinases in the PSC lysates were assessed with the Proteome Profiler Human Phospho-Kinase Array Kit (R&D system, CAT#: ARY003C), with 600 µg of PSC lysate being applied to the membrane. Membranes were imaged with the Bio-Rad ChemiDoc MP Imaging System.

### 2.16 PSC functional assay

PSCs were treated with exosomes freshly isolated from MIAPaCa-2 VC and NDRG1-expressing cells at 90% confluency. Cancer cell derived exosomes were suspended in IMDM media without FCS before treatment. To ensure an equal number of exosomes were given, 20% of VC-derived exosomes were removed (**Supplemental Figure 1)**, with the total number of exosomes for each PSC biological repeat being approx. 6.66 x 10^9^ particles per well in a 6 well plate. For MAPK activation analysis, PSCs were treated with exosomes for 0, 15, 30, 45, and 60 min in serum-free media to eliminate FBS contaminants. To assess PSC activation and collagen production, PSC cells were seeded on a 96-well plate with incubated with cancer cell derived exosomes for 24 h in serum-free media. Following exosome treatment, cell lysates were collected for protein quantification and immunoblotting. Additionally, PSCs were seeded in 96-well glass plates, incubated with cancer cell derived PHK67 stained exosomes for 24 h, following by the immunofluorescence protocol (described in *Section 2.8*). Fixed PSCs were examined for exosome uptake and collagen production with immunofluorescent microscopy. Images were taken with a Leica Stellaris 8 confocal microscope.

### 2.17 Exosome uptake assay

After isolation from PSCs or MIAPaCa-2 parental cells, exosomes were concentrated and suspended in 200 µL of exosome-free PBS. Exosomes were labelled with a PKH67 Green Fluorescent Cell Linker Kit (CAT#: PKH67GL-1KT, Sigma-Aldrich) following the manufacturer’s protocol. Labelled PSC-derived exosomes were then resuspended in serum free media and added to either MIAPaCa-2 VC or MIAPaCa-2 NDRG1 cells for 24 h. Prior to imaging, the MIAPaCa-2 cells were fixed with 4% PFA and washed with exosome free PBS to remove residual exosomes. Following DAPI staining and mounting, immunofluorescent imaging was performed to detect the PKH67-labelled PSC-derived exosomes using a Leica Stellaris 8 confocal microscope with a 488 nm laser.

### 2.18 Exosome-lysosome colocalization with live cell confocal microscopy

MIAPaCa-2 parental cell-derived exosomes were collected and stained with PKH67 dye as described above. MIAPaCa-2 VC and NDRG1 cells were seeded into black polystyrene microplates (CAT#: CLS3603, Sigma Aldrich) cultured under standard conditions. Upon reaching 75% confluency, they were treated with LysoTracker Deep Red (5nM) (CAT#: L1249. Themo Fisher Scientific) and Hoechst live nuclei stain (2 µg/mL) for l h at 37°C. FCS-free media containing PKH labelled exosomes (approx. 1 x 10^9^) was then added to either VC or NDRG1 cells, followed by a 2.5 h incubation. Confocal live-cell imaging was performed using a Leica stellaris 8 microscope for 16 h at 37°C and 5% CO2, capturing l6 ROIs/sample every 15 min (total 65 time points). Data analysis was performed using custom Java script to calculate the overlay score of lysosome and exosome fluorescent signals. Fluorescent signals were normalised to nuclear DAPI intensity.

### 2.19 Data Analysis

Densitometric data from Western blot analysis were evaluated using Bio-Rad Image Lab software. Protein expression levels were quantified as adjusted volumes and normalized to B-actin. Immunofluorescence intensities were measured in LAS X software, and colocalization scores (Pearson’s correlation coefficients) were calculated using the Fiji Coloc2 plugin. For the exosome–lysosome colocalization experiment, a custom JavaScript using Fuji was employed to batch-calculate the overlay scores between lysosomal and exosomal fluorescent signals. All graphs were generated in GraphPad Prism, and statistical significance was determined using Student’s t-test.

## 3. Results

### 3.1 NDRG1 reduces exosome release by interfering with intracellular trafficking and the ESCRT Pathway

NDRG1 was found to inhibit cross-talk between PaC cells and PSCs, disrupting key oncogenic signalling pathways, although the precise mechanisms underlying this inhibitory effect remain to be fully elucidated [30, 31, 60]. To investigate whether NDRG1 can influence exosome biogenesis in PaC, we generated a stable NDRG1-overexpressing MIAPaCa-2 cell line and compared it with a vector control (VC) MIAPaCa-2 line that exhibits low NDRG1 expression. Immunoblot analysis confirmed successful overexpression, with NDRG1 levels approximately 5-fold higher in the overexpressing cells compared to VC cells (**Figure 1B)**.

To isolate and characterize EVs derived from both NDRG1 overexpressing and VC MIAPaCa-2 cells, we used the well-established differential ultracentrifugation protocol to enrich for either: ***(i)*** large oncosomes following centrifugation at 2800 RCF (2.8K EVs); ***(ii)*** large EVs (microvesicles) following centrifugation at 10,000 RCF (10K EVs), and; ***(iii)*** small EVs (exosomes) following centrifugation at 100,000 RCF (100K EVs) [61]. Nanoparticle tracking analysis (NTA) revealed that NDRG1-overexpressing MIAPaCa-2 cells secreted significantly fewer exosomes than VC cells (**Figure 1C**). Quantitative analysis showed an approximately 25% reduction in particle number in the NDRG1-overexpressing condition (2.4×10^11^ *vs.* 3.2×10^11^ particles/ml). The size distribution profiles further confirmed this reduction across the characteristic exosome size range (30-200 nm), with NDRG1-overexpressing cells showing a lower peak and narrower distribution (**Figure 1D**). In addition, to further validate our EV isolation protocol, we also examined the size profile of the three subtypes of EVs using ZetaSizer Ultra (Malvern Panalytical, UK; **Table 2).** Each EV population was observed to have distinct particle sizes, with 2.8K EVs having an average size of 1240-1600 nm, 10K EVs being predominantly in the 500-600 nm range, and exosomes being approximately 200 nm. These findings were consistent with EV characterisation methods [62].

**Table 2:** UltraSizer report. Z-average size distribution and related statistical parameters of extracellular vesicles (EVs) isolated from MIAPaCa-2 cells, analysed using the UltraSizer system. Measurements were performed on three EV fractions (2.8K, l0K, and l00K) from both vector control (VC) and NDRG1-overexpressing cells. For each measurement, 10 μL of EV suspension was further diluted in 1 mL of ultrapure water. The table reports the mean particle size (Z-Average, in nm), standard deviation, relative standard deviation (RSD), and the minimum and maximum particle sizes detected.

| **Z-Average (nm)** | **Mean** | **Standard Deviation** | **RSD** | **Minimum** | **Maximum** |
| --- | --- | --- | --- | --- | --- |
| MIAPaCa-2 VC 2.8K | 572.4 | 166.6 | 29.11 | 428.5 | 754.9 |
| MIAPaCa-2 VC 10K | 575.6 | 98.83 | 17.17 | 497 | 686.5 |
| MIAPaCa-2 VC 100k | 203.4 | 9.332 | 4.589 | 195.6 | 213.7 |
| MIAPaCa-2 NDRG1 2.8K | 1615 | 614.5 | 38.05 | 905.6 | 1981 |
| MIAPaCa-2 NDRG1 10K | 572.4 | 166.6 | 29.11 | 428.5 | 754.9 |
| MIAPaCa-2 NDRG1 100K | 232.3 | 2.976 | 1.281 | 229.5 | 235.4 |

Transmission electron microscopy of the MIAPaCa-2 VC or NDRG1 derived exosome fractions confirmed the morphological characteristics of isolated exosomes, revealing the typical double-layered membrane structures with a diameter less than 200 nm from both conditions (**Figure 1E**). Notably, the ultrastructural features appeared consistent between VC and NDRG1 exosomes, suggesting that NDRG1 affects exosome quantity rather than basic morphology. Moreover, we performed immunoblot analyses for established EV markers across the different fractions derived from MIAPaCa-2 VC and NDRG1 cells (**Figure 1F**). As expected, the endosomal origin markers ALIX, TSG101, and the exosome membrane marker CD9 were predominantly enriched in the exosome fraction [63], with lower expression in the 2.8K and l0K fractions. Importantly, the mitochondrial marker, Mitofilin was detected primarily in the 2.8K and l0K fractions and was detected with minimal expression in the exosome fraction [64], confirming minimal cellular contamination in our exosome preparations. Comparing the VC and NDRG1 fractions, we observed reduced levels of ALIX and TSG101 in the exosome fraction from NDRG1-overexpressing cells compared to VC cells, suggesting that NDRG1 may reduce exosome production and affect the incorporation of these ESCRT components into exosomes.

To validate the effect of NDRG1 on exosome production in another PaC cell line, PANC-1 cells were also transfected to over-express NDRG1, which also led to a significant reduction in exosome secretion from PANC-1 NDRG1 cells compared to their VC counterparts (**Supplemental Figure 2 A–D)**. Conversely, NDRG1 knockdown in MIAPaCa-2 cells had the opposite effect, leading to a pronounced increase in exosome release (**Supplemental Figure 2E-G)**. Overall, these results validate the inhibitory effect of NDRG1 on exosome secretion by PaC cells.

To explore the underlying mechanism by which NDRG1 reduces exosome release, we first examined key ESCRT pathway proteins, including hepatocyte growth factor-regulated tyrosine kinase substrate (HRS), TSG101 and CD9, as well as the ESCRT accessory protein ALIX, which plays a central role in exosome biogenesis. Immunoblot analysis of cellular lysates from MIAPaCa-2 VC and NDRG1 cells revealed that overexpression of NDRG1 was associated with a significant reduction in ALIX and TSG101, while CD9 accumulated intracellularly and HRS remained unchanged (**Figure 1G**). A similar regulatory role for NDRG1 was also observed in the PANC-1 cells, where NDRG1 overexpression also led to decreased expression of ALIX and TSG101, while CD9 expression was upregulated (**Supplemental Figure 3A**). Notably, HRS, which initiates endosomal sorting [65], was significantly up-regulated by NDRG1 in the PANC-1 cells (**Supplemental Figure 3A**). Notably, ALIX was detected as 2-3 distinct bands in both MIAPaCa-2 and PANC-1 cells, which may arise from post-translational modifications[66, 67].

To further understand how NDRG1 affects intracellular endosome trafficking and MVB formation, the expression of small GTPases including Rab27a, Rab27b, Rab5a and Rab9a which are crucial for MVB maturation and membrane fusion [68, 69], and lysosomal marker LAMP-2 were also examined. Overexpression of NDRG1 in MIAPaCa-2 cells led to a significant reduction in Rab27a, Rab27b and Rab5a (**Figure 1H**). In contrast, Rab9a and LAMP-2 were upregulated in NDRG1 overexpressing MIAPaCa-2 cells (**Figure 1H**), suggesting enhanced lysosomal activity [70]. Similar results were observed in the PANC-1 cells, where Rab27A and Rab5a were decreased, while LAMP2 was increased in NDRG1 overexpressing cells (**Supplemental Figure 3B)**. However, Rab27b showed increased expression, while Rab9a was significantly reduced in NDRG1 overexpressing PANC-1 cells when compared to the VC cells (**Supplemental Figure 3B)**, which was opposite to the expression of these proteins in MIAPaCa-2 cells. These results suggest the effect of NDRG1 on exosome biogenesis may involve slightly different mechanisms in the 2 PaC cell lines examined.

Knocking down NDRG1 (NDRG1*^KD^*) in MIAPaCa-2 cells had no significant effect on the ESCRT proteins, although there was a significant increase in Rab27a expression when compared to the negative control (NC) cells (**Supplemental Figure 3C, D**). It is important to note that in these latter experiments, NDRG1 was not completely silenced, and this may contribute to the lack of effect on the other proteins examined.

Overall, these findings indicate that NDRG1 overexpression significantly reduces exosome secretion, potentially by altering key components of the ESCRT pathway and modulating intracellular trafficking machinery in PaC cells. The concurrent downregulation of Rab GTPases (Rab27a/b, Rab5a) which promote exosome secretion, and upregulation of lysosome-associated proteins (LAMP2, Rab9a) suggests that NDRG1 may redirect MVB content toward lysosomal degradation rather than exosomal release. Based on these observations, we next sought to determine whether NDRG1 functionally interacts with specific ESCRT factors to regulate exosome biogenesis.

### 3.2 NDRG1 interacts with ALIX and induces its proteasomal degradation

To determine whether NDRG1 functionally associates with the ESCRT pathway, we next examined its relationship with ALIX, a component of the ESCRT machinery that facilitates exosome maturation and cargo loading during MVB sorting [71]. Given the importance of ALIX in forming intraluminal vesicles (ILV), any modulation of ALIX expression or function has the potential to significantly impact exosome-mediated TME communication [29, 72].

Immunofluorescence microscopy performed on MIAPaCa-2 cells showed that NDRG1 overexpression markedly reduced ALIX levels in the cytoplasm, when compared with VC cells (**Figure 2A**), aligning with our earlier immunoblot analysis (**Figure 1G, H**). In addition, NDRG1 and ALIX were found to co-localize, as demonstrated by the Pearson’s correlation coefficient, which was substantially higher in the NDRG1-overexpressing cells (**Figure 2B**), suggesting a close spatial relationship between the two proteins. NDRG1 was also found to co-localize with Rab5a and Rab9a, as demonstrated by the significantly increased Pearson’s correlation coefficient between NDRG1 and each of these proteins in MIAPaCa-2 cells (**Supplemental Figure 4A - D**).

**Figure 2.**
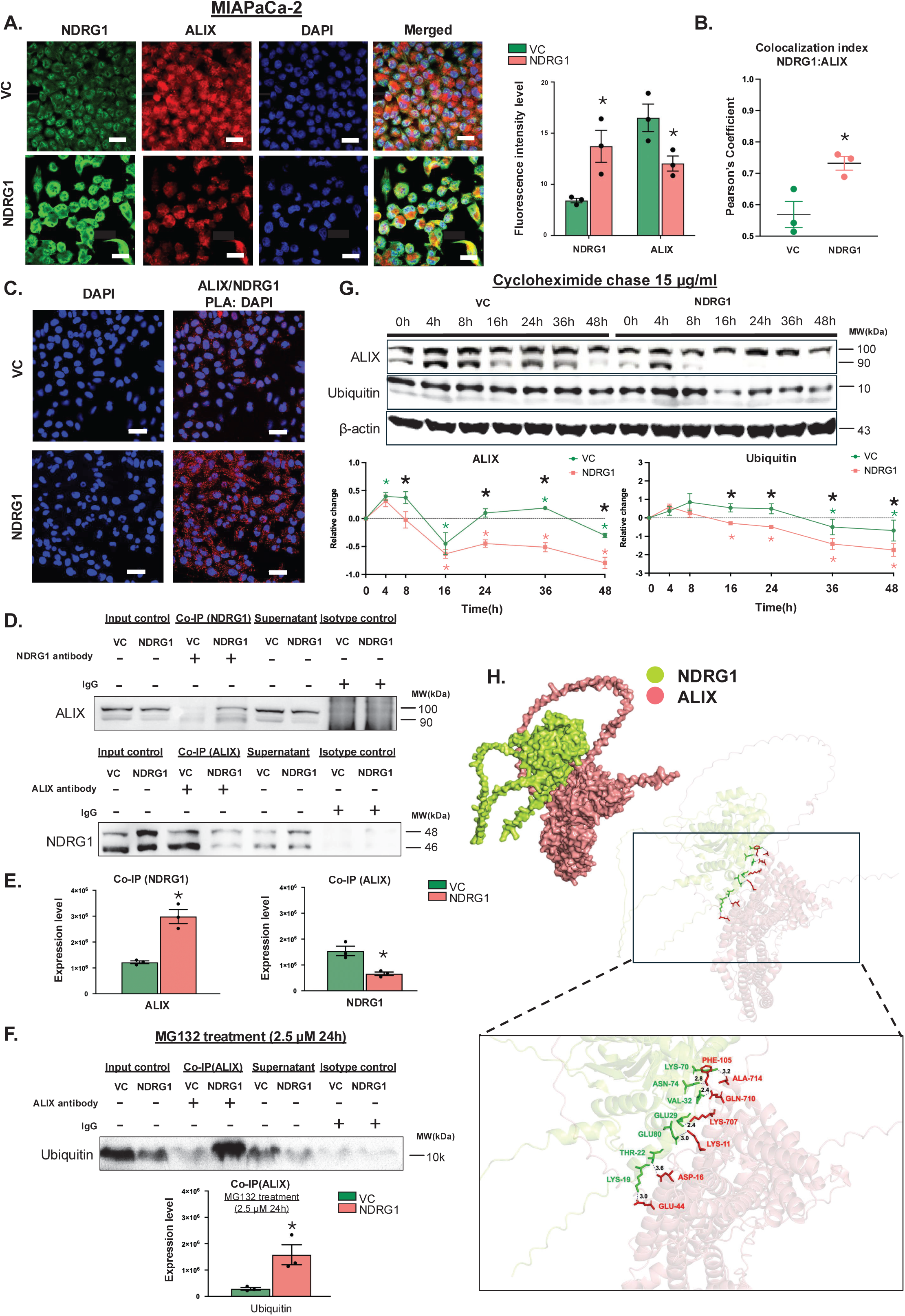
NDRG1 interacts with ALIX and promotes its proteasomal degradation. (**A)** Representative immunofluorescence images of MIAPaCa-2 cells expressing vector control (VC) or NDRG1 show staining for NDRG1 (green), ALIX (red), and nuclei (DAPI, blue), with merged images indicating colocalization of NDRG1 and ALIX (Scale bar: 20 μm). Quantification of fluorescence intensity levels are shown in the adjacent bar graph, with each dot representing an independent experiment and bars indicating mean ± SEM. (**B**) Pearson’s correlation coefficient analysis quantifies the degree of colocalization between NDRG1 and ALIX. (**C**) Proximity ligation assay (PLA) images reveal protein-protein interactions between NDRG1 and ALIX (red), with DAPI staining nuclei (blue) (Scale bar: 20 μm). (**D**) Co-immunoprecipitation (Co-IP) experiments confirm the interaction between NDRG1 and ALIX, with ALIX detected in NDRG1 pull-downs and *vice versa*. (**E**) Quantification of Co-IP data shows levels of ALIX-NDRG1 interaction, represented as mean ± SEM from 3 independent experiments. (**F)** Increased ALIX ubiquitination is observed in NDRG1-expressing cells following MG132 treatment (2.5 μM, 24 h) and quantification of Co-IP data shows ALIX ubiquitination, represented as mean ± SEM from 3 independent experiments. (**G)** Representative western blot images demonstrate a cycloheximide (CHX) chase of ALIX and ubiquitin protein levels in VC and NDRG1 cells over a 48-h time-course following CHX treatment (l5 µg/mL), with B-actin as a loading control. Coloured asterisks (*) indicate significant differences compared to 0 h, while black asterisks indicate significant difference between VC and NDRG1 cells. Quantification of ALIX and ubiquitin levels at each time point was normalized to B-actin. All Statistical significance was determined using Student’s *t*-test: \**p* < 0.05. (**H**) AlphaFold3 protein-protein interaction prediction of ALIX (pink) and NDRG1 (green). Enlarged view shows the binding amino acid residues from NDRG1 (green) and ALIX (red). Dashed lines indicate polar interactions labelled with distance between binding amino acids (Å3).

Given that ALIX directly interacts with ESCRT-III and promotes MVB maturation [71], this observation raised the possibility that NDRG1 might influence exosome composition by directly altering ALIX’s availability and function. To investigate the potential interaction between NDRG1 and ALIX further and assess direct protein–protein interaction, a proximity ligation assay (PLA), which detects proteins residing within close molecular distances (:S40 nm) was used. The PLA signals indicated a notable increase in the NDRG1–ALIX interaction in cells overexpressing NDRG1 (**Figure 2C**), with only minimal background signal being observed from control samples where only one antibody (NDRG1 or ALIX) or samples with no antibody were imaged (**Supplemental Figure 4E**). This was further corroborated by co-immunoprecipitation assays, showing that ALIX was detected in the NDRG1 immunoprecipitate and *vice versa* (**Figure 2D**), with significantly higher levels of ALIX bound to NDRG1 in NDRG1-overexpressing cells compared with VC cells (**Figure 2E**). Notably, the amount of NDRG1 protein detected in the ALIX immunoprecipitate was markedly lower in the NDRG1 overexpressing lysates (**Figure 2E**), likely reflecting the overall reduced levels of ALIX in these cells. Together with the Western blot data from **Figure 1G** and the immunofluorescence data from **Figure 2A**, this further suggests that ALIX might be subjected to more rapid turnover in the NDRG1 overexpressing MIAPaCa-2 cells.

To assess if the NDRG1-mediated reduction of ALIX could be linked to lysosomal degradation, we examined the potential co-localization of ALIX with the lysosomal marker LAMP-2 in MIAPaCa-2 VC and NDRG1 cells **(Supplemental Figure 4F)**. However, we observed no marked increase in co-localization of these 2 proteins when comparing VC to NDRG1 overexpressing cells **(Supplemental Figure 4F)**. We next investigated the involvement of proteasomal degradation by treating MIAPaCa-2 VC and NDRG1 cells with the proteasomal inhibitor MG132 for 24 h to inhibit proteasomal turn-over of ubiquitinated proteins and thus enable detection of protein ubiquitination by Co-IP. Treatment of cells with MG132 followed by immunoprecipitation of ALIX revealed increased levels of ubiquitin binding to ALIX in the NDRG1-overexpressing cells (**Figure 2F**), implicating NDRG1 as a regulator that targets ALIX for proteasomal degradation.

This finding was further substantiated by cycloheximide (CHX) chase experiments, which measure protein turnover by inhibiting new protein synthesis. In VC cells, ALIX levels declined noticeably at 16 h then increased at 24 and 36 h, followed by a reduction at 48 h CHX post-treatment. However, in NDRG1-overexpressing cells, significant ALIX depletion was evident as early as 16 h and remained significantly lower than in VC cells for all subsequent time-points (**Figure 2G)**. Additionally, the rate of ubiquitin turnover was elevated in the NDRG1-overexpressing cells, suggesting that increased proteasomal activity accompanies or is driven by the presence of high NDRG1 levels (**Figure 2G**).

Finally, to gain insight into how NDRG1 might physically interact with ALIX and potentially modulate its function in exosome biogenesis, we performed *in silico* protein–protein docking using UniProt sequences and AlphaFold3 for molecular interaction [58]. These simulations indicated that NDRG1 is binding to multiple regions of the V-shaped Bro1 domain of ALIX (**Figure 2H**). The Bro1 domain of ALIX encompasses amino acid 1 to amino acid 359 [73]. NDRG1 was shown to bind to ALIX amino acid sequence LYS-11, ASP-16, GLU-44 and PHE-105. Notably, the Bro1 domain mediates ALIX’s interaction with CHMP4B which belongs to ESCRT-III [71, 74], and is essential for MVB maturation and cargo loading [29, 71]. We also noted that engagement by NDRG1 altered ALIX’s conformational state (**Supplemental Figure 5A**), which could potentially restrict its access to binding partners (**Supplemental Figure 5B, C**). Hence, NDRG1 binding to ALIX may have a dual role including enhancing the proteasomal degradation of ALIX, while also potentially impairing ALIX-mediated cargo recruitment and MVB formation. Overall, these data demonstrate that NDRG1 binding to ALIX has significant implications for exosome composition and release in PaC.

### 3.3 NDRG1 Expression Alters EV Protein Cargo

Building on our observation that NDRG1 modulates ALIX in the ESCRT pathway, we performed comprehensive proteomic analysis to determine whether these alterations translate into measurable changes in EV cargo. Using label-free quantitative mass spectrometry, we compared the protein composition of EVs derived from NDRG1-overexpressing MIAPaCa-2 cells with those from VC cells.

Examining exosomes, our analysis revealed a total of 2,906 proteins across both conditions, with 2,331 proteins detected in exosomes from both cell types. A striking asymmetry emerged in the distribution of unique proteins, with VC exosomes containing 575 unique proteins, whereas NDRG1 exosomes contained only 96 unique proteins. This marked reduction in protein diversity suggests that NDRG1 significantly restricts the range of cargo packaged into exosomes (**Figure 3A**).

**Figure 3.**
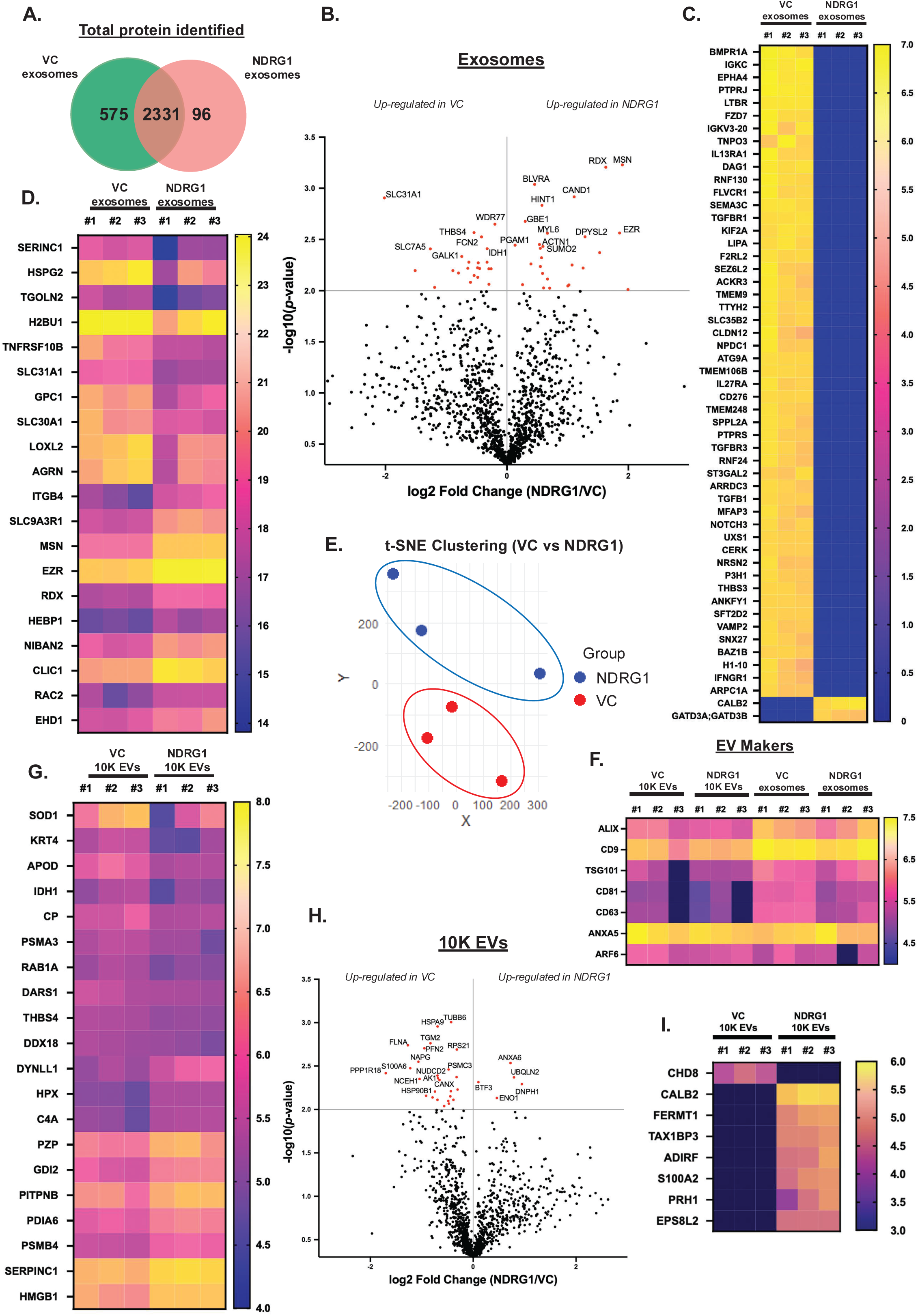
Mass spectrometry proteomic analysis reveals NDRG1 expression affects peptide cargo in EVs. (**A**) Venn diagram illustrating the number of proteins identified in exosomes from MIAPaCa-2 cells expressing VC or NDRG1. (**B**) Volcano plot showing the log2 fold change (NDRG1/VC) on the x-axis and −log10 (*p*-value) on the y-axis, representing differentially expressed proteins in exosomes isolated from MIAPaCa-2 cells. Red dots indicate proteins with significant differential expression (*p* < 0.01). (**C**) Heatmap showing the top 50 proteins uniquely and consistently expressed proteins in VC exosomes, ranked based on average expression levels, compared to NDRG1 exosomes ranked by fold change. (**D**) Heatmap showing the top 20 differentially expressed proteins in exosomes isolated from MIAPaCa-2 cells expressing VC or NDRG1, identified using proteomic mass spectrometry with a library-free search approach. The first 10 proteins (top rows) are more highly expressed in VC exosomes, while proteins ranked 11–20 (bottom rows) are more highly expressed in NDRG1 exosomes. (**E**) t-Distributed Stochastic Neighbour Embedding (t-SNE) plot showing clustering of proteomic profiles from exosomes isolated from MIAPaCa-2 cells expressing VC or NDRG1. Each point represents an independent biological replicate, with VC exosomes (red) and NDRG1 exosomes (blue). (**F**) Heatmap showing the expression levels of EV markers (ALIX, CD9, TSG101, CD81, CD63, ANXA5, and ARF6) in 10K EVs and exosomes isolated from MIAPaCa-2 cells expressing VC or NDRG1. (**G**) Heatmap showing the top 20 differentially expressed protein cargos in 10K EVs isolated from MIAPaCa-2 cells expressing VC or NDRG1. The first 10 proteins (top rows) are more highly expressed in VC 10K EVs, while proteins ranked 11–20 (bottom rows) are more highly expressed in NDRG1 10K EVs. **(H**) Volcano plot representing differentially expressed proteins in 10K EVs isolated from MIAPaCa-2 cells. Proteins significantly upregulated in VC 10K EVs are shown on the left, while those upregulated in NDRG1 10K EVs are shown on the right. Red dots indicate proteins with significant differential expression (*p* < 0.01, Statistical significance was determined using Student’s *t*-test). (**I**) Heatmap displaying proteins uniquely and consistently expressed in 10K EVs from MIAPaCa-2 cells expressing VC or NDRG1. In all heatmaps, each column represents an independent biological replicate (#1, #2, #3). All data were normalized and log10-transformed, with yellow indicating higher expression and blue indicating lower expression and black indicating no expression. Statistical significance was determined using Student’s *t*-test.

Statistical analysis identified distinct protein signatures that distinguished NDRG1 exosomes from their VC counterparts. Transport-related proteins such as SLC31A1 (copper transporter) and amino acid transporter SLC7A5, along with metabolic enzymes including phosphoglycerate mutase 1 (PGAM1) [75], isocitrate dehydrogenase 1 (IDH1) [76], and galactokinase 1 (GALK1) [77], were significantly enriched in VC exosomes (**Figure 3B**). In contrast, NDRG1 exosomes showed elevated levels of cytoskeletal regulatory proteins including moesin (MSN), ezrin (EZR), and cadherin-related protein cullin-associated and neddylation-dissociated 1 (CAND1), as well as signalling molecules such as biliverdin reductase A (BLVRA) and hit protein 1 (HINT1) (**Figure 3B**).

A heatmap of differentially and uniquely expressed proteins in VC exosomes (**Figure 3C**) highlighted factors involved in TGF-B signalling (TGFBl, TGFBRl, BMPRlA) [78], immune modulation (CD276, IGKV3-20, ILl3RAl, LTBR) [79-8l], matrix remodelling (MFAP3, THBS3), and intracellular trafficking/autophagy (ATG9A, SNX27, SPPL2A, VAMP2) [82–85]. Iron metabolism related heme export and cholesterol/lipid metabolic regulators (FLVCR1, LIPA) [86, 87] and signal transduction mediators (PTPRJ, PTPRS [88]) were also enriched in VC exosomes. Further, NOTCH3, which is one of the most studied CAF activating factors [89], was only found in VC exosomes, indicating that VC-derived exosomes carry distinct functional cargo compared to those from NDRG1-overexpressing cells. In contrast, only 2 proteins were found to be consistently and uniquely expressed in all 3 biological replicates of exosomes derived from NDRG1 overexpressing cells, including calcium binding protein CALB2 and mitochondrial proteins GATD3A/B (**Figure 3C**).

Among the most differentially expressed proteins, the membrane protein SERINC1 showed the highest upregulation in VC exosomes, followed by HSPG2, TGOLN2, H2BU1, and TNFRSF10B. Notably, the glycoprotein GPC1, previously reported as a PaC exosome marker [90, 91], was significantly enriched in VC exosomes. In contrast, cytoskeletal regulators MSN, EZR, and RDX demonstrated substantially higher abundance in NDRG1 exosomes across all replicates [92] (**Figure 3D**). Dimensionality reduction analysis using t-SNE demonstrated clear separation between VC and NDRG1 exosome proteomes (**Figure 3E**), confirming that NDRG1 expression substantially reshapes the overall exosomal protein landscape.

To confirm the identity and purity of our EV preparations, we also examined the distribution of canonical EV markers across both 10K EV and exosome fractions (**Figure 3F**). As expected, exosome membrane tetraspanins (CD9, CD8l, CD63) and ESCRT-associated proteins (ALIX, TSG101; [93]) were predominantly detected in the exosome fractions, while the 10K EV fractions showed higher expression of EV markers such as ANXA5 and ARF6 [94, 95].

Interestingly, NDRG1’s influence on protein cargo extended beyond exosomes to 10K EV as well. In the 10K EV fraction, we observed distinct protein signatures that differentiated EVs derived from VC *vs.* NDRG1 overexpressing cells. Among the top 10 up-regulated genes in VC 10K EVs were the antioxidant enzyme SOD1, structural protein KRT4, lipid carrier APOD, metabolic enzyme IDH1, and ceruloplasmin (CP). Conversely, the top 10 up-regulated genes in NDRG1 10K EVs included hemopexin (HPX), complement component C4A, pregnancy zone protein (PZP), and GDP dissociation inhibitor 2 (GDI2). This distinct pattern of differential protein expression was consistent across all three biological replicates (**Figure 3G**).

Additional significant alterations in 10K EV cargo included upregulation of mitochondrial tubulins (TUBB8), filamin A (FLNA), transglutaminase 2 (TGM2), and interferon regulatory factor (IFRH2) in VC 10K EVs, while annexin A6 (ANXA6), ubiquilin-2 (UBQLN2), basic transcription factor 3 (BTF3), and 2’-deoxynucleoside 5’-phosphate N-hydrolase 1 (DNPH1) were elevated in NDRG1 10K EVs **(Figure 3H)**. Finally, we also observed that NDRG1 10K EVs had some exclusively expressed proteins that were not detected in VC 10K EVs **(Figure 3I)**, including calcium binding proteins, calbindin 2 (CALB2) and S100 calcium binding protein A2 (S100A2), cytoskeleton remodelling proteins and cell adhesion proteins, such as EPS8 Like 2 (EPS8L2) and fermitin family member 1 (FEMT1). In contrast, the chromatin remodelling protein CHD8 was exclusively expressed in the VC l0K EVs (**Figure 3I**).

Collectively, our comprehensive proteome analysis demonstrates that NDRG1 expression substantially alters the protein cargo profiles of both exosomes and 10K EVs in MIAPaCa-2 cells. These proteomic changes involve proteins associated with multiple cellular processes, including signalling pathways, metabolism, cytoskeletal organization, and vesicular trafficking, suggesting that NDRG1 broadly influences the composition of extracellular vesicles secreted by PaC cells.

## 4. Exosomes derived from NDRG1 overexpressing PaC cells attenuate PSC activation

The proteomic analysis above revealed that TGFBl signalling components were significantly downregulated in exosomes from NDRG1-overexpressing cancer cells (**Figure 3C**). Immunoblot analysis confirmed markedly reduced TGFBl expression in NDRGl exosomes compared to VC exosomes, with this reduction being consistent across all EV fractions (2.8K, 10K, and 100K) derived from NDRG1-overexpressing cells (**Figure 4A**). Both precursor (55kDa) and mature (l2kDa) forms of TGFBl showed lower abundance in NDRGl-derived exosomes. Given the key role TGFB1 plays in driving PSC activation and ECM deposition in the TME [96, 97], we investigated the functional consequences of this altered exosomal cargo on PSC behaviour by incubating PSCs with exosomes isolated from either VC or NDRG1-overexpressing MIAPaCa-2 cancer cells (**Figure 4B**).

**Figure 4.**
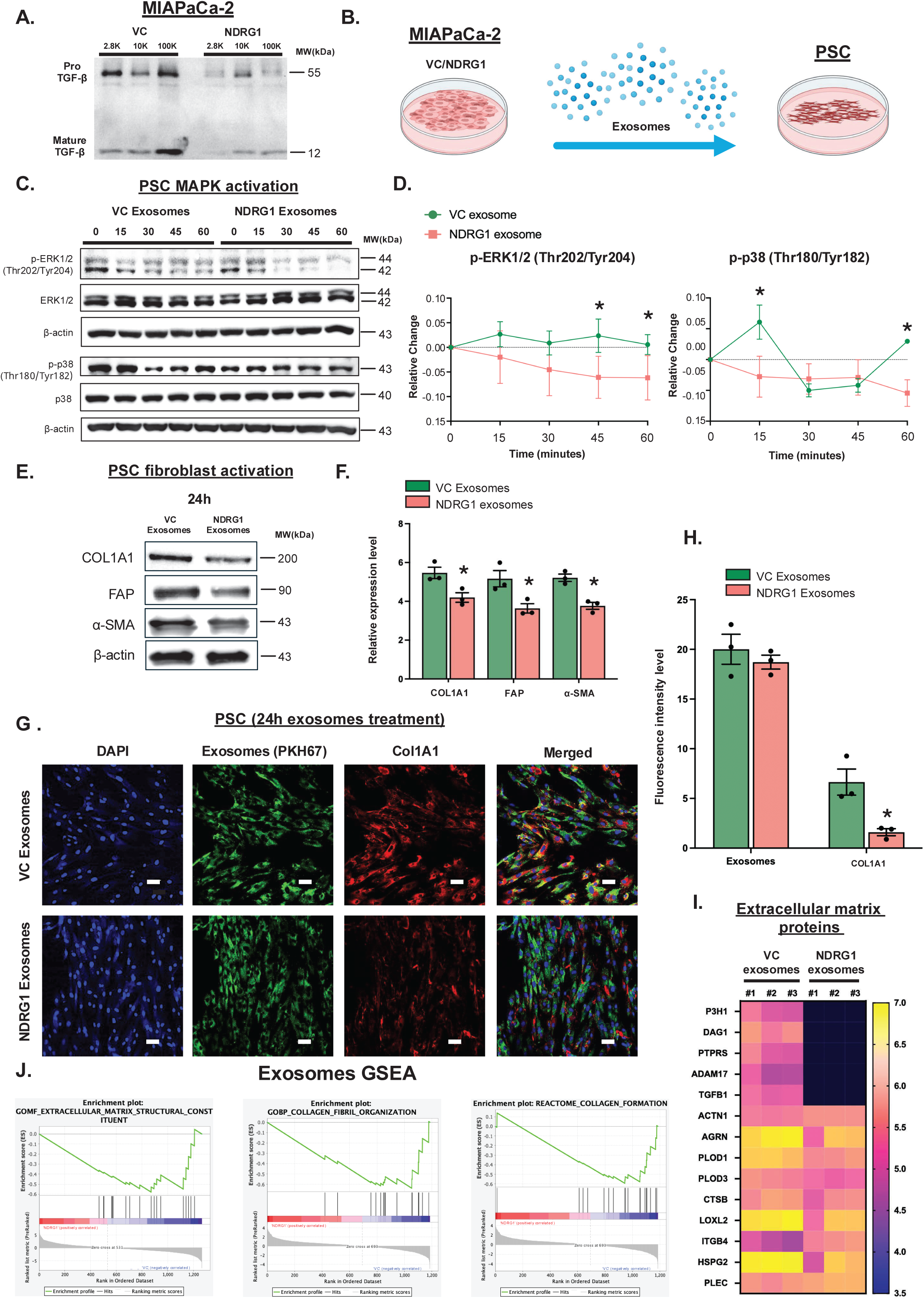
NDRG1 exosomes reduce PSC activation and collagen production compared to VC exosomes. (**A**) Representative western blot images showing the levels of pro-TGF-B and mature TGF-B in EV fractions (2.8K, 10K, and 100K) isolated from VC or NDRG1 MIAPaCa-2 cells. (**B**) Schematic illustrating the experimental setup for exosome treatment. (**C**) Representative western blot images of MAPK pathway activation in PSCs treated with equal amounts of exosomes from VC or NDRG1-expressing MIAPaCa-2 cells for the indicated time points (0, 15, 30, 45, 60 min). Levels of phosphorylated ERK1/2 (p-ERK1/2) and P38 (p-p38) were assessed relative to total ERK1/2 and p38, respectively, with B-actin serving as a loading control. (**D**) Quantification of relative changes in p-ERK1/2 and p-p38 phosphorylation over time, normalized to total ERK1/2 and P38, respectively. Asterisks (*) indicate significant expression differences between VC and NDRG1 exosome treated PSCs. (**E**) Representative western blot images of fibroblast activation markers (COL1A1, FAP and a-SMA) in PSCs treated with equal amounts of exosomes from VC or NDRG1-expressing MIAPaCa-2 cells for 24 h, and their densitometric analysis (**F**), with B-actin serving as a loading control. (**G**) Representative immunofluorescence images of PSCs treated with exosomes from VC or NDRG1-expressing cells for 24 h. Nuclei were stained with DAPI (blue), exosomes are labelled with PKH67 (green), and COL1A1 (collagen type I, red) marks PSC collagen production. Scale bar: 40 μm. (**H**) The bar graph quantifies fluorescence intensity levels of exosome uptake (PKH67) and COL1A1 expression from **G**. (**I**) Heatmap showing the significantly differentially expressed extracellular matrix (ECM) proteins in exosomes isolated from MIAPaCa-2 cells expressing VC or NDRG1. Each column represents an independent biological replicate (#1, #2, #3). Data were normalized and log10-transformed, with yellow indicating higher expression and blue indicating lower expression and black indicating no expression. (**J**) Gene set enrichment analysis (GSEA) plots illustrate the enrichment of ECM-related pathways in exosomes isolated from MIAPaCa-2 cells expressing VC or NDRG1, based on proteomic mass spectrometry data. The analysed pathways include GOMF_Extracellular Matrix Structural Constituent, GOBP_Collagen Fibril Organization, and REACTOME_Collagen Formation. In all bar graphs, each dot represents an independent experiment, with bars indicating mean ± SEM. Statistical significance was determined using Student’s *t*-test and is indicated as \**p* < 0.05. To normalize for differences in exosome secretion, exosome input was adjusted based on yield quantification, with VC exosome volume reduced by 20% to match the lower secretion levels of NDRG1-derived exosomes (See **Supplemental** Figure 1).

To investigate the signalling alterations in PSCs following exosome-mediated communication, we first utilized a human phospho-kinase array kit to examine the phosphorylation status of key kinases. PSCs were treated with equal amounts of exosomes derived from either VC or NDRG1-overexpressing MIAPaCa-2 PaC cells. Our analysis showed a notable difference in kinase activation profiles between the two treatment groups. Specifically, the phosphorylation of several critical components of the MAPK signalling pathway, a key regulator of cellular proliferation, differentiation, and stress responses [98, 99] were found to be significantly downregulated in PSCs treated with exosomes from NDRG1-overexpressing cancer cells (**Supplemental Figure 6A, B**). These proteins include ERKl/2 and p38a, which have previously been implicated to induce activation of PSCs into CAFs [100–102]. Time-course analysis of PSCs exposed to cancer cell-derived exosomes revealed distinct phosphorylation patterns of ERKl/2 (Thr202/Tyr204) and p38a (Thrl80/Tyrl82) between exosome treatment groups (**Figure 4C, D**). PSCs treated with exosomes derived from NDRG1 overexpressing cells exhibited significantly reduced ERKl/2 phosphorylation at 45 and 60 min compared to those treated with VC exosomes (**Figure 4C, D**). Similarly, p38a phosphorylation was significantly lower in PSCs treated with exosomes derived from NDRG1 overexpressing cells at 15 and 60 min timepoints (**Figure 4C, D**). Total ERKl/2 and p38a levels remained unchanged throughout the incubation period confirming that the observed differences reflected altered activation rather than protein abundance.

We next examined whether this attenuated MAPK signalling affected PSC activation into CAFs. PSCs exposed to exosomes derived from NDRG1 overexpressing MIAPaCa-2 cells had significantly lower levels of classical CAF markers, including COL1A1, FAP and a-SMA, when compared to PSCs exposed to VC-derived exosomes (**Figure 4E, F)**. These findings are consistent with our previous observations that conditioned media from NDRG1-overexpressing cancer cells maintains PSCs in a more quiescent state [30].

To validate these findings at the cellular level, we performed immunofluorescence microscopy on PSCs incubated with fluorescently labelled (PKH67) exosomes from either VC or NDRG1-overexpressing MIAPaCa-2 cells for 24 h (**Figure G**). Exosomes from both VC and NDRG1 overexpressing cancer cells were taken up by PSCs at similar rates, as demonstrated by the green fluorescence from the PKH67 dye accumulating in PSCs (**Figure 4G**). However, the expression of COL1A1 (Collagen type I alpha 1; red signal) was dramatically reduced in PSCs exposed to NDRG1 exosomes. Quantification of fluorescence intensity confirmed significant collagen reduction, with approximately 80% lower COLlA1 expression in NDRG1 exosome-treated PSCs compared to the VC group (**Figure 4H**). As exosome uptake itself was not significantly different between conditions, this suggests that the observed effects stemmed from altered exosomal cargo rather than differential uptake efficiency.

Given the decrease in collagen production, we next examined whether this reflected broader changes in the extracellular-matrix cargo of NDRG1-derived exosomes. Heatmap analysis revealed differential expression of multiple ECM-associated proteins between NDRG1 and VC exosomes across all three biological replicates (**Figure 4I**). Several key ECM components, including prolyl 3-hydroxylase (P3H1) [103], dystroglycan 1 (DAG1 [104]), protein tyrosine phosphatase receptor type S (PTPRS), and ADAM metallopeptidase 17 (ADAM17), were not detected in NDRG1 exosomes. Moreover, peptides such as Agrin (AGRN) [105], procollagen-lysine,2-oxoglutarate 5-dioxygenase 1 (PLOD1) [106], procollagen-lysine,2-oxoglutarate 5-dioxygenase 3 (PLOD3), cathepsin B (CTSB), lysyl oxidase-like 2 (LOXL2) [107] [l08], and heparan sulfate proteoglycan 2 (HSPG2) exhibited significantly higher expression levels in VC derived exosomes compared to exosomes derived from NDRG1-overexpressing cancer cells. Gene Set Enrichment Analysis (GSEA) of the proteomic data provided further evidence of functional divergence between exosome populations. Pathways related to ECM organization and collagen synthesis, specifically “ECM structural constituent,” “collagen fibril organization,” and “collagen formation", were significantly enriched in VC exosomes (**Figure 4J**), with pronounced enrichment scores and distinct clustering of ECM-related genes.

Together, these results demonstrate that exosomes from NDRG1-overexpressing cancer cells disrupt TGFBl-mediated MAPK activation and substantially reduce CAF marker expression and collagen production in PSCs, highlighting NDRG1’s potential role in modulating tumour-stroma interactions through altered exosomal communication.

## 5. NDRG1 reduces exosome uptake and promotes exosome–lysosome interactions

To extend our understanding of how NDRGl modulates cross-talk between PaC cells and the TME, we also investigated whether NDRG1 influences cancer cell uptake of exosomes. Exosomes can serve as nutrient sources under metabolic stress, and previous evidence suggests PaC cells may “rewire” nearby CAFs to supply metabolite-rich exosomes [109]. Our earlier observations (**Figure 1G**) showed that Rab5a, a key protein regulating early endosomal trafficking [110], was downregulated in NDRGl-overexpressing cancer cells. This prompted us to examine whether NDRG1 can also influence exosome uptake by PaC cells.

Exosomes were enriched from PSC conditioned media, with NTA and immunoblotting confirming particle size and marker expression was consistent with exosomes (**Figure 5 A, B**). Exosomes were stained with PKH67 dye and their concentration normalized to ensure consistent exosome amounts were loaded onto both MIAPaCa-2 VC and NDRG1 overexpressing cells, followed by an overnight incubation (**Figure 5C**). Confocal immunofluorescence revealed significantly fewer PKH67-labelled exosomes were detected in NDRGl-overexpressing cells compared with VC cells (**Figure 5D, E**). This was further validated using PANC-1 cells, where the NDRG1 overexpressing PANC-1 cells internalized significantly fewer PSC-derived exosomes (**Supplemental Figure 6C)**.

**Figure 5.**
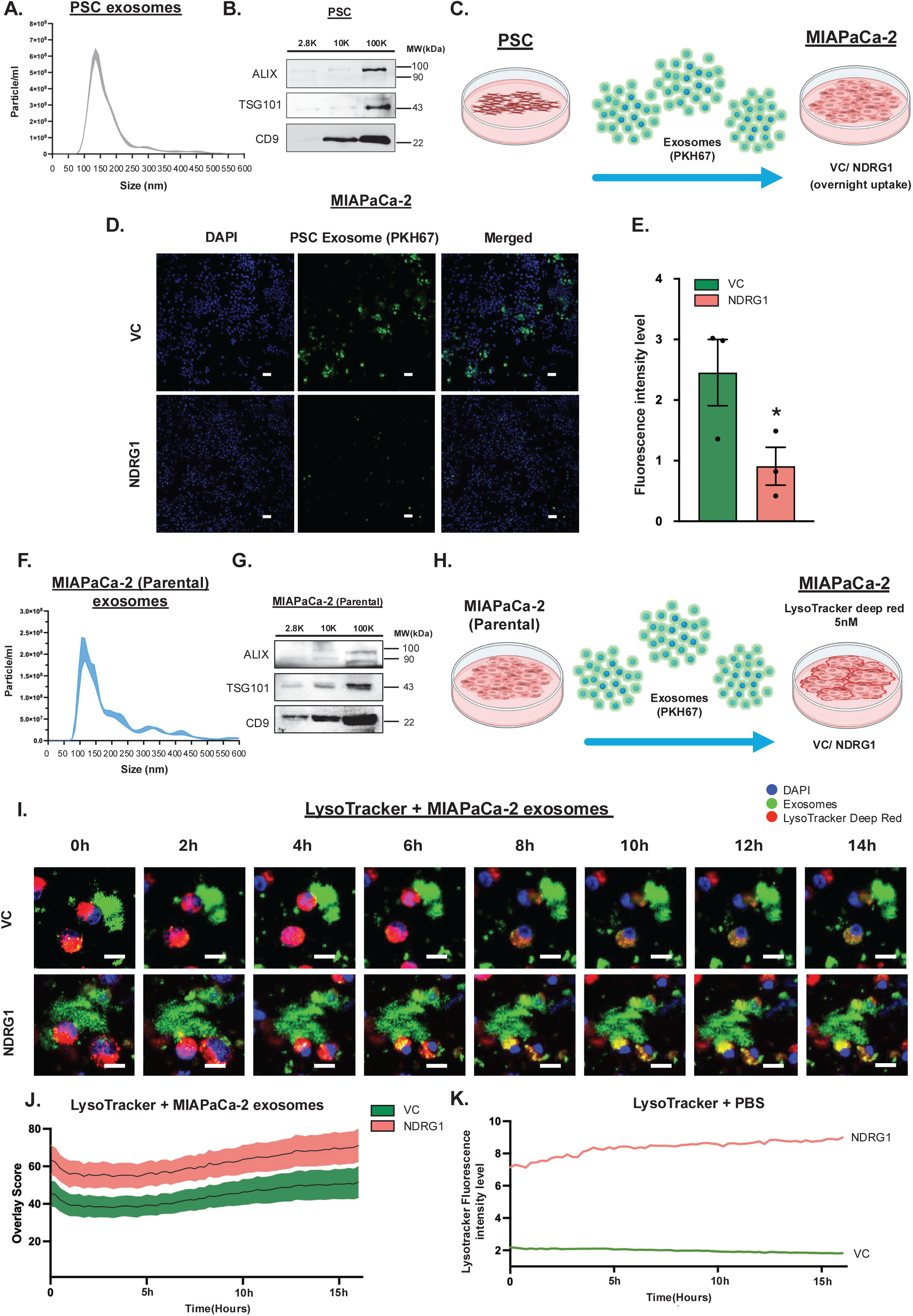
NDRG1 decreases exosome uptake from the TME and directs internalized exosomes to the lysosome. **(A)** NTA showing the size distribution of PSC-derived 100K exosomes. (**B**) Representative western blot images of exosome markers (ALIX, TSG101, and CD9) in PSC-derived EVs confirming the presence of exosomal markers in the 100K fraction. (**C**) Schematic illustrating the experimental setup where pancreatic stellate cells (PSCs) secrete exosomes, which are labelled with PKH67 (green) and subsequently incubated overnight with MIAPaCa-2 cells expressing VC or NDRG1. (**D**) Representative immunofluorescence images of VC or NDRG1-expressing MIAPaCa-2 cells incubated with PSC-derived exosomes overnight. Nuclei are stained with DAPI (blue), and PKH67-labeled PSC exosomes are green. Merged images indicate exosome uptake. Scale bar: 60 μm. (**E**) Quantification of exosome uptake fluorescence intensity in NDRG1-expressing MIAPaCa-2 cells compared to VC cells. Each dot represents an independent experiment, with bars indicating mean ± SEM. Statistical significance was determined using Student’s *t*-test, * *p* < 0.05. (**F**) NTA showing the size distribution of MIAPaCa-2-derived 100K exosomes. (**G**) Representative western blot images of exosome markers (ALIX, TSG101, and CD9) in MIAPaCa-2-derived EVs confirming the presence of exosomal markers in the 100K fraction. (**H**) Schematic illustrating the experimental setup where exosomes from MIAPaCa-2 parental cells (PKH67-labeled, green) were added to VC and NDRG1-expressing MIAPaCa-2 cells pre-treated with LysoTracker Deep Red (5 nM) and Hoechst live nuclei stain (2 μg/mL) for 1 h, which is followed by a 2.5 h incubation with exosomes to ensure uptake prior to imaging. (**I**) Representative time-lapse confocal microscopy images showing exosome uptake and trafficking in VC and NDRG1-expressing MIAPaCa-2 cells over time. DAPI (blue) stains nuclei, PKH67 (green) marks exosomes, and LysoTracker Deep Red (red) labels lysosomes. Confocal live-cell imaging was performed with Leica stellaris 8 for l6 hours at 37°C and 5% CO2, capturing l6 ROIs/sample every l5 min (total 65 time points). Yellow colour indicate colocalization of exosomes with lysosomes. Scale bar: 20 μm. (**J**) Quantification of colocalization (overlap score) between exosomes and lysosomes in VC and NDRG1-expressing cells, calculated using the Fiji Multiply function. The overlap score represents the degree of exosome-lysosome colocalization over time. The shaded area represents the standard SEM at every timepoint. (**K**) Quantification of LysoTracker fluorescence intensity over time in control conditions (PBS-treated cells). Fluorescence intensity and overlay score were normalized to the intensity of the nucleus.

We next explored whether NDRG1 also modulates the intracellular fate of internalized exosomes. Our earlier findings (**Figure 1G**) and previous research [111, 112], indicate that NDRGl can regulate lysosomal function, raising the possibility that TME-derived exosomes might be redirected for lysosomal degradation in NDRGl-overexpressing cells. To investigate this, we isolated and validated exosomes from parental MIAPaCa-2 cells **(Figure 5F, G)**, followed by labelling with PKH67 dye. MIAPaCa-2 VC or NDRG1 cells were pre-incubated with lysosomal stain LysoTracker Deep Red, followed by incubation with the PKH67-labelled exosomes (**Figure 5H)**. Live-cell confocal imaging was then performed over l6 h under standard cell culture conditions to track whether internalized exosomes were co-localizing with lysosomes (**Figure 5I)**.

Colocalization measurements confirmed a significantly greater overlap between PKH-labelled exosomes (green) and lysosomes (red) in the NDRG1 overexpressing cells, as demonstrated by the increased abundance of yellow puncta (**Figure 5I**) and reflected by the higher overlay score of the two markers (**Figure 5J**). Analysis of the confocal images also indicated significant increase in basal lysosomal activity over the course of 16 h in NDRGl-overexpressing MIAPaCa-2 cells relative to VC cells (**Figure 5K)**, consistent with our earlier observations of elevated LAMP2 in NDRG1 overexpressing PaC cells (**Figure 1H**). These data suggest that NDRG1 not only reduces the number of exosomes taken up by cancer cells, but also directs them towards the lysosomes, potentially limiting the metabolite or signalling advantages these exosomes confer within the TME.

## 6. Discussion

It has been well established that the TME is a key contributor to PaC progression and development of resistance to current therapies. However, targeting the TME in PaC has proven challenging due to its dynamic nature that is constantly being reshaped by both cancer and stromal cells. This is mediated by the bi-directional cross talk between cancer and stromal cells, which exchange EVs, growth factors, metabolites and cytokines to reprogram the function of neighbouring cells and manifest a favourable niche for PaC cells to proliferate and metastasise [113, 114]. In the current study, we demonstrate for the first time that the metastasis suppressor NDRG1 can inhibit the cellular communication between PaC cells and the TME, with this effect being mediated by disruption of exosomes release, biogenesis, cargo packaging and uptake by cancer cells. NDRG1 significantly attenuated exosome release from PaC cells, an effect that was likely due to the ability of NDRG1 to inhibit exosome biogenesis machinery by potentially interacting with and regulating the expression of various ESCRT proteins, including, HRS, TSG101, ALIX, and CD9.

NDRG1 reduced the expression of the ESCRT-0 protein TSG101, which is involved in ESCRT initiation, maturation of early and late endosomes and MVB cargo sorting [115]. This suggests that NDRG1 expression can inhibit the initiation of exosome biogenesis potentially by regulating the ESCRT-0 complex.

ALIX is another important ESCRT pathway regulator and a key component of ESCRT-III that is predominantly involved in late endosomal and MVB cargo selection [116-ll8]. We demonstrate that NDRG1 down-regulates ALIX expression through ubiquitin-mediated proteasomal degradation. Studies indicate that ALIX facilitates exosome cargo sorting, secretion, and drives malignant phenotypes in transformed cancer cells [117, 119]. Furthermore, ALIX-dependent exosome secretion has been linked to tumour immune evasion and metastasis, as it is involved in packaging immune-regulatory proteins such as PD-L1 into exosomes, thereby modulating anti-tumour immunity [72]. A number of earlier studies have shown that NDRG1 can regulate the expression of other proteins by facilitating their ubiquitination and subsequent proteasomal degradation in pancreatic and other cancers [120–122]. This suggests that NDRG1 has a broader functional effect on the proteasomal degradation machinery, a hypothesis that is further supported by the increased turnover of ubiquitin observed in the NDRG1 overexpressing PaC cells in this study.

Notably, besides promoting ALIX ubiquitination and eventual degradation, the direct binding of NDRG1 to ALIX may also inhibit its function in the first instance. ALIX and its interaction with ESCRT-III, is crucial for ILV formation and protein sorting during MVB maturation [123, 124]. The predicted spatial interaction between NDRG1 and ALIX, particularly at the Bro1 domain which mediates ESCRT-III recruitment, suggests that NDRG1 may interfere with critical protein-protein interactions required for MVB formation. This mechanism differs from previously described regulators of exosome biogenesis that typically act through transcriptional control of ESCRT components or modulation of endosomal trafficking without direct protein-protein interactions [125].

In addition to ESCRT pathway proteins, we also found that CD9, a surface marker for exosomes, was significantly upregulated in PaC cell lines by NDRG1. Interestingly, CD9 upregulation was previously related to increased lysosome number, while deletion of CD9 promoted MVB formation and exosome secretion in melanoma cancer cells [126]. These findings demonstrate that the ESCRT complex is intimately associated with endo-lysosomal vesicular structure formation and maturation, and that NDRG1 might influence both intracellular processes.

This was further confirmed by our observations that NDRG1 altered the expression of major regulators of endo-lysosomal activities, namely the Rab GTPase family members Rab5a, Rab27a and Rab9a. Work by Ostrowski *et al.* identified that reduced levels of either Rab27a/b, or Rab5a can lead to decreased exosome secretion [23, 68]. However, these different Rab GTPases oversee different parts of intracellular vesicular transportation. For instance, Rab27a is the “docking-and-fusion” specialist positioned at the cell periphery, while Rab27b shepherds membranes from the trans-Golgi network (TGN) toward MVEs, priming them for subsequent Rab27a-dependent fusion [68]. Rab5a is known as a key endosomal trafficking regulator that mainly controls endocytosis and early endosome formation [127–129]. Hence, our observation that NDRG1 overexpression reduced cancer cell exosome secretion and uptake, can also be explained by impaired MVB maturation and failed initiation of endocytosis due to the alterations in these Rab GTPases.

In agreement with our observations, others have also shown that NDRG1 may regulate EV biogenesis through its interactions with vesicular trafficking machinery. NDRG1 can directly interact with Prenylated Rab Acceptor 1 protein (PRA1), which then regulates small GTPase Rab proteins that facilitate transport between cell organelles [38]. Moreover, evidence suggests that NDRG1 directly binds to Rab4a to regulate E-cadherin transport and recycling through an endosome-related mechanism [33]. In addition, NDRG1-silenced cells were found to have impaired lysosomal function and increased secretion of neutral sphingolipids and ceramides [32, 34, 49], which are known to affect exosome biogenesis [130]. This is consistent with our findings in MIAPaCa-2 cells where NDRG1 overexpression resulted in increased levels of lysosomal marker LAMP-2 and Rab9a, which is involved in lysosome formation [70, 131]. Interestingly, there is evidence suggesting that NDRG1 also regulates lysosome and proteasome activities [111, 112, 122]. Ortega *et al.* discovered that NDRG1 silencing increased exosome release in breast, colon and ovarian cancer cell lines, with this effect being accompanied by inhibition of lysosomal acidification [112]. The abovementioned effects of NDRG1 on lysosomal degradation of numerous proteins may potentially be mediated by its effects on or association with Rab9a.

Another important finding of this study is the alteration of exosomal protein cargo in NDRG1-overexpressing cells. Our study reveals that NDRG1 expression fundamentally reshapes the EV proteome in PaC cells, selectively restricting the packaging of pro-tumorigenic, metabolic, and stromal-activating proteins. Through comprehensive proteomic analysis of both exosomes and 10K EVs, we show that NDRG1 narrows the range of vesicular cargo and actively modulates intercellular communication in a manner consistent with its well-established role as a metastasis suppressor [35, 132, 133]. One of the most striking findings was the reduction in the diversity of exosomal proteins secreted by NDRG1-overexpressing cells compared to VC. The loss of over 80% of unique exosomal cargo highlights NDRGl’s capacity to globally restrict the dissemination of signalling molecules through exosomes. This reduction likely limits the tumour cells’ ability to interact with and modify the TME, representing a novel mechanism by which NDRG1 enforces a less aggressive cellular phenotype.

Importantly, the exclusion of proteins involved in TGF-B signalling—including TGFBl, TGFBR1, and BMPR1A—from NDRG1-derived exosomes suggests a specific suppression of tumour-stroma crosstalk pathways. TGF-B family members are known to activate PSCs, promoting the dense desmoplastic stroma that characterizes pancreatic tumours and fuels metastasis and therapy resistance [134, 135]. In addition, VC-derived exosomes were enriched in NOTCH3, a key regulator implicated in the activation of CAFs, which can further exacerbate fibrosis, immune suppression, and tumour progression [136]. The absence of TGF-B and NOTCH3 signalling components in NDRG1 exosomes strongly supports a model in which NDRG1 limits stromal activation and remodelling through selective exosome cargo exclusion.

An additional and critical layer of EV cargo reprogramming by NDRG1 involves metabolic crosstalk. Exosomes derived from VC cells were enriched in key metabolic enzymes such as IDH1, PGAM1, GALK1, as well as nutrient transporters including SLC31A1 and SLC7A5. These metabolic enzymes and transporters are critical for tumour metabolism, supporting processes such as redox balance, glycolysis, biosynthesis, nutrient uptake, and mTORC1-driven metabolic reprogramming essential for cancer growth and adaptation [137–141]. Together, these proteins enable metabolic reprogramming of the TME by supplying biosynthetic intermediates, enhancing nutrient uptake, and reshaping redox balance in recipient stromal and immune cells [142]. The selective exclusion of these metabolic regulators from NDRG1 EVs suggests inhibition of tumour-promoting metabolic symbiosis, further supporting the metastasis-suppressor role of NDRG1.

Interestingly, NDRG1 exosomes selectively incorporated BLVRA, an enzyme intimately involved in heme catabolism, antioxidant defence, and iron metabolism [143, 144]. BLVRA activity reduces biliverdin to bilirubin, a potent antioxidant that mitigates reactive oxygen species (ROS) accumulation, and its activity is tied to maintaining iron homeostasis [143–146]. This observation is especially significant given that NDRG1 is known to be transcriptionally upregulated in response to iron chelation and modulates intracellular iron levels [30, l47, l48]. The selective inclusion of BLVRA in EVs suggests that NDRG1 may regulate redox and iron stress not only within tumour cells but also in neighbouring stromal and immune cells through vesicle-mediated signalling.

Building upon the proteomic analysis, which revealed significant changes in protein cargo from NDRG1 overexpressing PaC cells, we further demonstrate that these molecular alterations have functional consequences on PSCs. Notably, phosphorylation of p38 and pERKl/2, both downstream effectors of the MAPK pathway, was reduced in PSCs following exposure to exosomes from NDRG1 overexpressing PaC cells. This was accompanied by reduced expression of CAF markers and reduced collagen production, suggesting that NDRG1 expression in PaC cells reduces activation of PSCs into CAFs. This is consistent with our previous reports that NDRG1 overexpression in PaC cells maintains PSCs in a quiescent, non-fibrogenic state [149], and demonstrates that exosomes are an important messenger for this stromal reprogramming.

The functional impact on ECM remodelling was further reinforced by our proteomic profiling of EV cargo. Several ECM-related proteins critical for collagen crosslinking and matrix remodelling, such as P3H1, DAG1, PTPRS, and ADAM17, were absent in NDRG1-derived exosomes. Moreover, multiple ECM remodelling enzymes, including LOXL2, PLOD1, PLOD3, CTSB, and HSPG2, were significantly more abundant in VC-derived exosomes. GSEA supported these observations, showing enrichment of pathways related to collagen fibril organization and ECM structural constituents exclusively in VC exosomes. These data suggest that NDRG1 expression not only reduces the delivery of soluble profibrotic signals but also limits the transmission of structural ECM components that would otherwise support desmoplasia. These findings further underscore NDRG1’s role as a key regulator of tumour– stroma interactions, acting through the strategic modulation of extracellular vesicle communication to reshape the TME toward a less fibrotic and less supportive niche for tumour progression.

In addition, our finding that NDRG1 not only affects cancer cell-derived exosomes but also modulates the uptake and processing of PSC-derived exosomes is particularly significant. Recent studies have suggested that cancer cells may “educate” stromal cells to provide metabolite-rich exosomes that support tumour growth under metabolic stress [60, 109]. In the current study we demonstrate that NDRG1 not only limits the uptake of PSC-derived exosomes by cancer cells but also directs internalized exosomes toward lysosomal degradation, effectively constraining the pro-tumorigenic advantages conferred by intercellular vesicle transfer. NDRG1’s ability to reduce exosome uptake and promote their lysosomal degradation suggests it may block this metabolic support system, potentially contributing to its tumour-suppressive function.

## 7. Limitations and Future Directions

While our study provides compelling evidence for NDRG1’s role in regulating exosome biology in PaC, several questions remain. The precise mechanism by which NDRG1 selectively alters exosomal cargo warrants further investigation, particularly regarding the enrichment of certain proteins amid general cargo restriction. Additionally, the effect of NDRG1-mediated exosome regulation on immune cell interactions within the tumour microenvironment remains to be explored.

## 8. Conclusion

In conclusion, our study establishes NDRG1 as a critical regulator of exosome biogenesis and intercellular communication in PaC. By interfering with the ESCRT pathway through direct interaction with ALIX, NDRG1 restricts exosome production and reshapes exosomal cargo to disrupt oncogenic signalling networks. These results highlight a novel extracellular dimension to NDRG1’s metastasis-suppressor role and suggest that modulating EV cargo selection could represent a new therapeutic strategy in PaC.

