## Supplemental Figures for "Silencing the signal: The metastasis suppressor NDRG1 disrupts exosome-mediated crosstalk in pancreatic cancer"

**Supplemental Figure 1:** NanoSight report comparing MIAPaCa-2 vector control (VC) and NDRG overexpressing (NDRG1) cell derived exosome concentration when VC exosomes were reduced by 20%. Exosome normalization ensures equal particle input for PSC treatment. NTA report showing exosome quantification after adjusting VC-derived exosome input. VC and NDRG1 exosomes particle concentrations after adjustment are shown in bold.

**Supplemental Figure 2:** (**A**) Western blot analysis showing NDRG1 expression in PANC-1 vector control (VC) cells and PANC-1 cells stably transfected to over-express NDRG1 (NDRG1), with β-actin as a loading control. The accompanying densitometry analysis quantifies relative NDRG1 expression, with each dot representing an independent experiment. (**B**) Nanoparticle tracking analysis (NTA) showing the concentration of small EVs (exosomes) isolated from VC or NDRG1 cells with each dot representing an independent experiment (bar graph). The size distribution graph depicts the particle size profile of exosomes measured by NTA, with shaded regions representing the standard error of the mean (SEM), calculated from three independent experiments. (**C**) Representative transmission electron microscopy (TEM) images of small EVs isolated from PANC-1 VC and NDRG1 cells. Scale bar: 200 nm. (**D**) Representative western blot images of exosomal markers (ALIX, TSG101, and CD9) and mitochondrial marker (Mitofilin) in 2.8K, 10K, and 100K EV fractions isolated from PANC-1 VC and NDRG1 cells. (**E**) Western blot analysis showing NDRG1 expression in MIAPaCa-2 negative control (NC) cells and MIAPaCa-2 cells stably transfected to knock-down NDRG1 ($NDRG1^{KD}$), with β-actin as a loading control. The accompanying densitometry analysis quantifies relative NDRG1 expression, with each dot representing an independent experiment. (**F**) NTA showing the concentration of small EVs (exosomes) isolated from NC or $NDRG1^{KD}$ cells with each dot representing an independent experiment (bar graph). The size distribution graph depicts the particle size profile of exosomes measured by NTA, with shaded regions representing the SEM, calculated from three independent experiments. (**G**) Representative western blot images of exosomal markers (ALIX, TSG101, and CD9). Each dot represents an independent experiment, and bars indicate the mean ± SEM. Statistical significance was determined by Student's *t*-test and is indicated as $*p < 0.05$; $***p < 0.001$.

**Supplemental Figure 3:** (**A**) Representative western blot images of ESCRT pathway proteins (HRS, ALIX, TSG101, and CD9) and their densitometric analysis in VC or NDRG1 PANC-1 cells. (**B**) Representative western blot images of endosomal trafficking regulators (Rab27a, Rab27b, Rab5a, Rab9a, and LAMP-2) and their densitometric analysis in VC or NDRG1 PANC-1 cells. (**C**) Representative western blot images of ESCRT pathway proteins (ALIX, TSG101, and CD9) and their densitometric analysis in negative control (NC) or NDRG1 knockdown ($NDRG1^{KD}$) MIAPaCa-2 cells. (**D**) Representative western blot images of endosomal trafficking regulators (Rab27a, Rab5a and LAMP-2) and their densitometric analysis in NC or $NDRG1^{KD}$ MIAPaCa-2 cells. For all densitometric analysis, band intensity was normalized to β-actin. Each dot represents an independent experiment, and bars indicate the mean ± SEM. Statistical significance was determined by Student's *t*-test and is indicated as $*p < 0.05$; $***p < 0.001$.

**Supplemental Figure 4:** (**A**) Representative immunofluorescence images of MIAPaCa-2 cells expressing vector control (VC) or NDRG1, showing staining for NDRG1 (green), Rab9A (red), and nuclei (DAPI, blue), with merged images indicating colocalization of NDRG1 and Rab9A. Scale bar: 20 µm. Quantification of fluorescence intensity levels for NDRG1 and Rab9A in VC and NDRG1 expressing cells is shown in the adjacent bar graph. (**B**) Pearson's correlation coefficient analysis quantifies the degree of colocalization between NDRG1 and Rab9A. (**C**) Representative immunofluorescence images of MIAPaCa-2 cells expressing VC or NDRG1 showing staining for NDRG1 (red), Rab5A (green), and nuclei (DAPI, blue), with merged images indicating colocalization of NDRG1 and Rab5A (Scale bar: 20 µm). Quantification of fluorescence intensity levels for NDRG1 and Rab5A in VC and NDRG1 expressing cells is shown in the adjacent bar graph. (**D**) Pearson's correlation coefficient analysis quantifies the degree of colocalization between NDRG1 and Rab5A. (**E**) Control proximity ligation assay (PLA) images of VC and NDRG1 MIAPaCa-2 cells incubated with no antibody, ALIX antibody (ab) alone and NDRG1 ab alone, with DAPI staining for nuclei (Scale bar: 20 µm). (**F**) Representative immunofluorescence images of MIAPaCa-2 cells expressing VC or NDRG1 showing staining for ALIX (red), LAMP-2 (green), and nuclei (DAPI, blue), with merged images. Scale bar: 20 µm. (**G**) Pearson's correlation coefficient analysis quantifies the degree of colocalization between ALIX and LAMP-2. In all bar graphs, each dot represents an independent experiment, with bars indicating mean ± SEM. Statistical significance was determined using Student's *t*-test, * $p < 0.05$.

**Supplemental Figure 5: AlphaFold3 Prediction of ALIX and NDRG1 interaction.** (**A**) ALIX show conformational changes before (blue) and after NDRG1 binding (pink). (**B**) Side view showing NDRG1 binding to the Bro-1 domain of ALIX. (**C**) Binding site between NDRG1 and ALIX shown as surface, while the entire proteins are shown as cartoon.

**Supplemental Figure 6:** (**A**) Proteome Profiler Human Phospho-Kinase Array (R&D Systems, Cat# ARY003C) was used to assess the phosphorylation status of key kinases in PSCs treated with exosomes from VC or NDRG1-expressing MIAPaCa-2 cells (**B**) Enlarged representative phospho-kinase array blots from **A** show differential phosphorylation patterns of ERK1/2 (T202/Y204, T185/Y187) and p38α (T180/Y182) in PSCs treated with VC or NDRG1 exosomes. (**C**) Representative immunofluorescence images of VC or NDRG1-expressing PANC-1 cells incubated with PSC-derived exosomes overnight. Nuclei are stained with DAPI (blue), and PKH67-labeled PSC exosomes (green). Merged images indicate exosome uptake. Scale bars represent 60 µm. Quantification of exosome uptake fluorescence intensity in NDRG1-expressing PANC-1 cells compared to VC cells is shown in the adjacent bar graph. Each dot represents an independent experiment, with bars indicating mean ± SEM. Statistical significance was determined using Student's *t*-test, * $p < 0.05$.

### MIAPaCa-2 VC Exosome

NANOSIGHT

Capture 2025-02-13 10-50-54

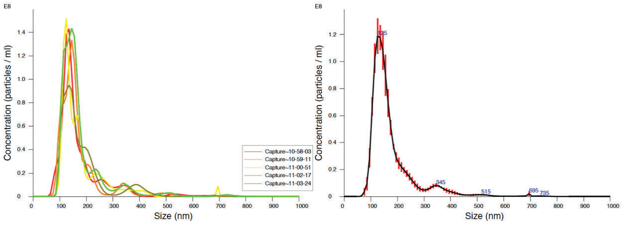


#### Stats: Mean +/- Standard Error

| **Metric** | **Value** |
|---|---|
| Mean | 175.7 ± 1.7 nm |
| Mode | 136.0 ± 4.3 nm |
| SD | 89.5 ± 1.3 nm |
| D10 | 98.8 ± 1.6 nm |
| D50 | 137.8 ± 1.7 nm |
| D90 | 266.0 ± 6.8 nm |
| **Concentration** | **1.06e+009 ± 5.30e+007 particles/ml** |
| Particles/frame | 54.0 ± 2.7 |

### MIAPaCa-2 NDRG1 Exosome

NANOSIGHT

Capture 2025-02-13 11-30-07

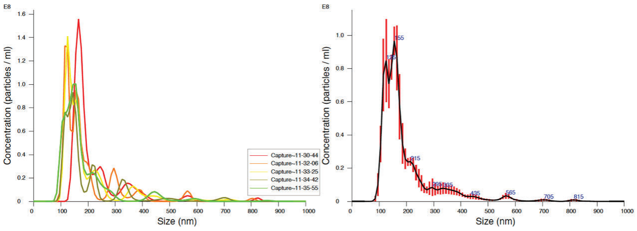


#### Stats: Mean +/- Standard Error

| **Metric** | **Value** |
|---|---|
| Mean | 197.9 ± 6.3 nm |
| Mode | 141.0 ± 8.3 nm |
| SD | 110.4 ± 2.7 nm |
| D10 | 110.6 ± 7.7 nm |
| D50 | 148.4 ± 5.3 nm |
| D90 | 319.8 ± 15.6 nm |
| **Concentration** | **9.27e+08 ± 4.99e+07 particles/ml** |
| Particles/frame | 47.0 ± 2.5 |

**Supplemental Figure 1**

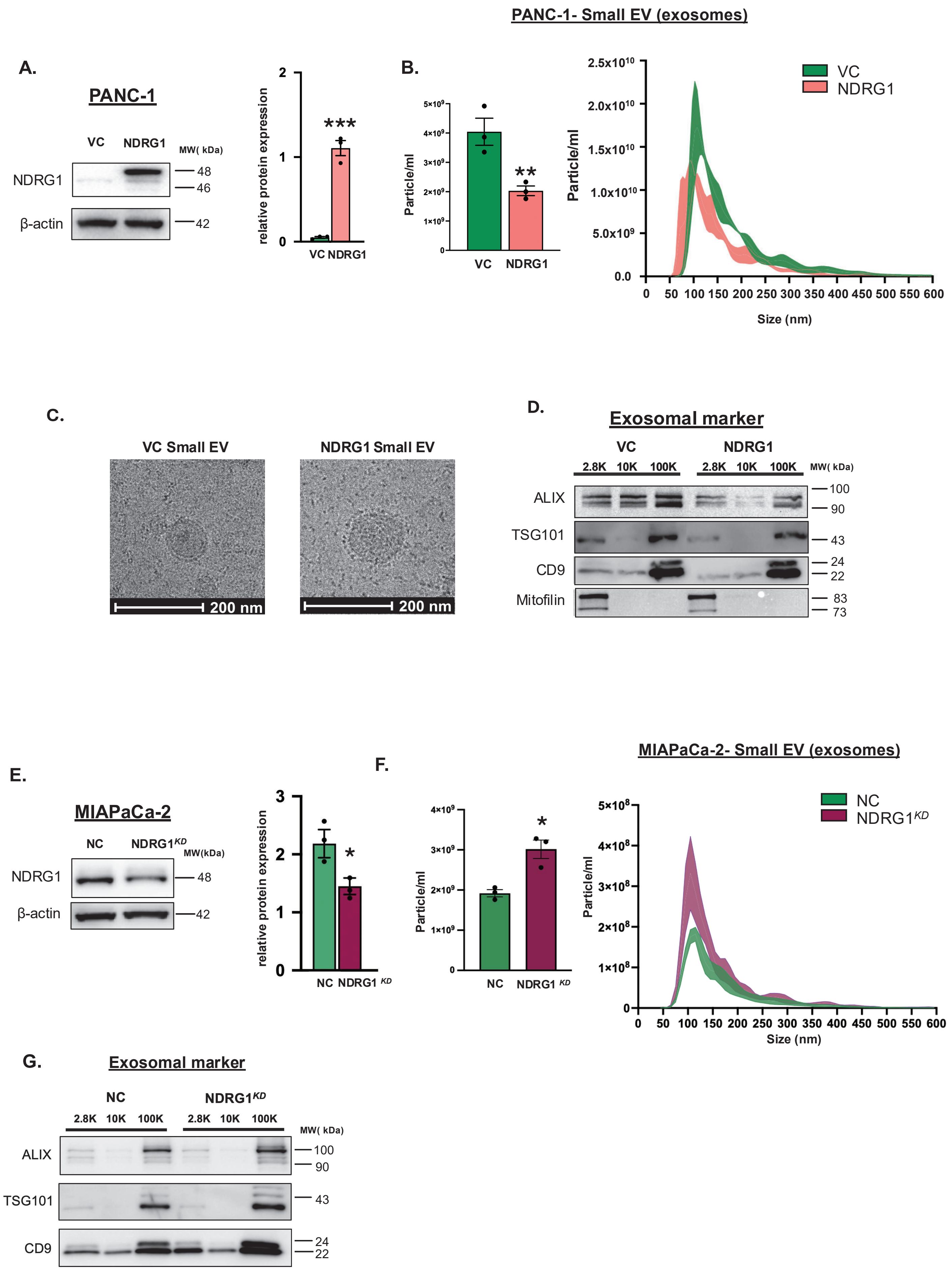


Supplemental Figure 2

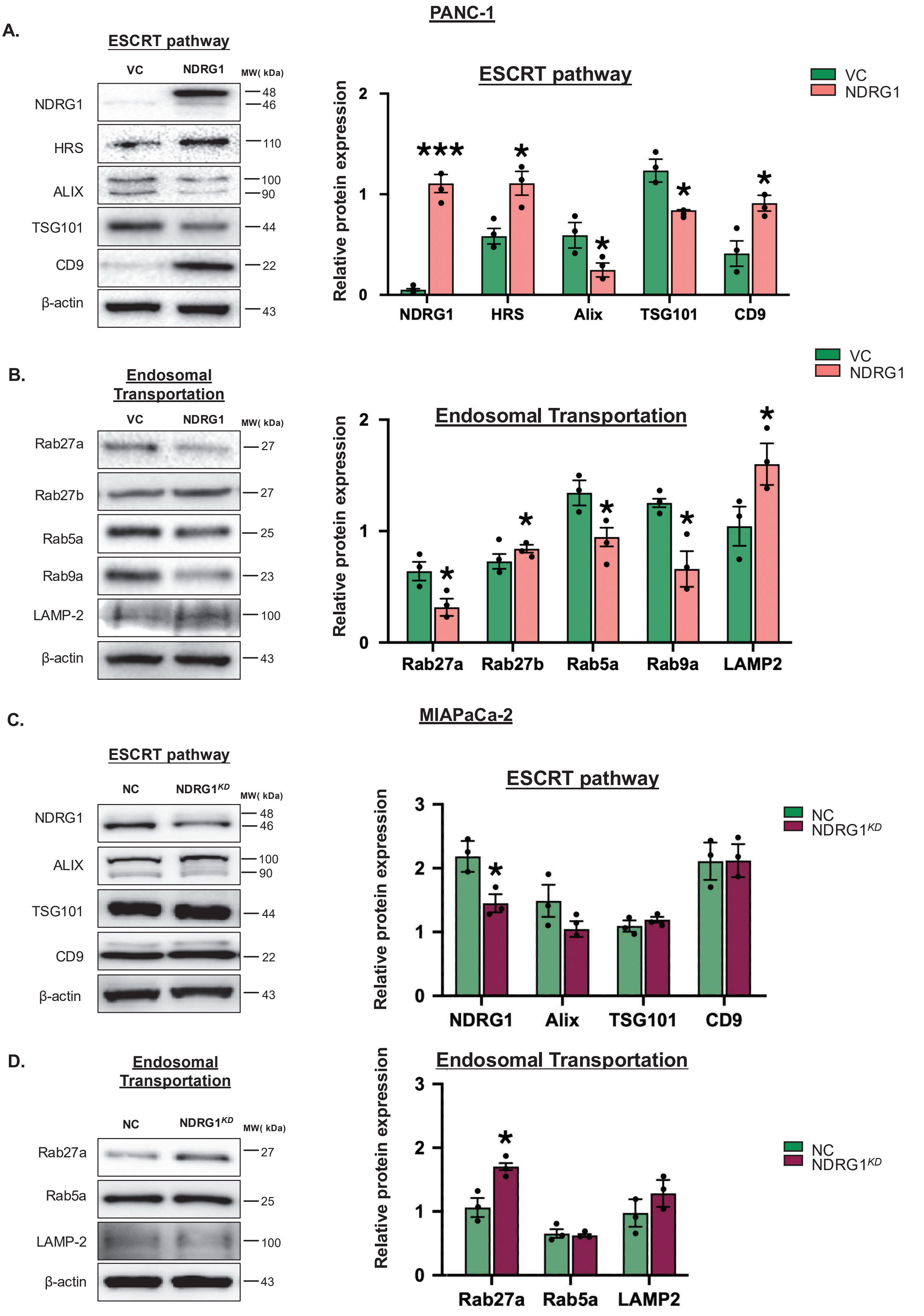


**Supplemental Figure 3**

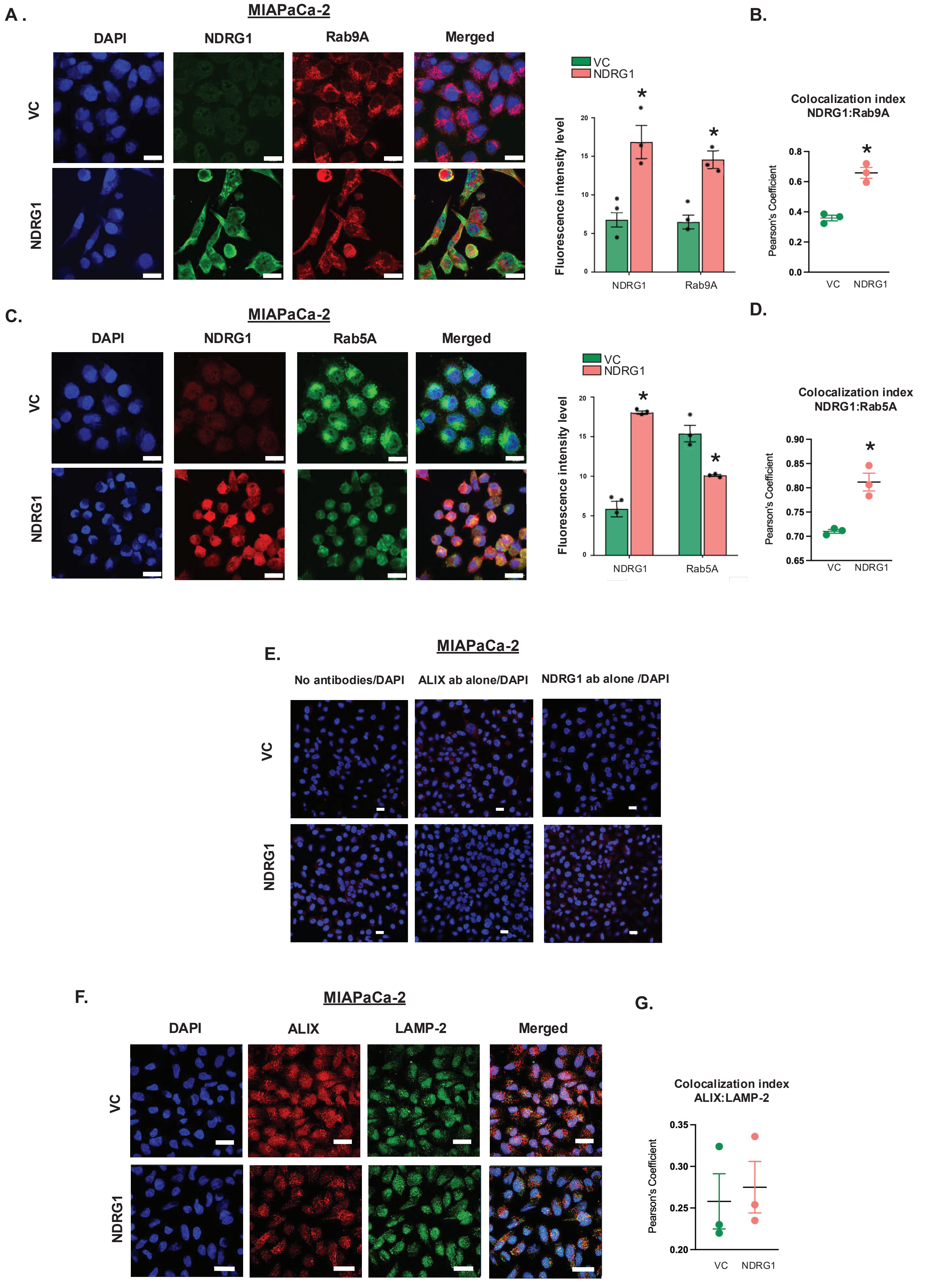


Supplemental Figure 4

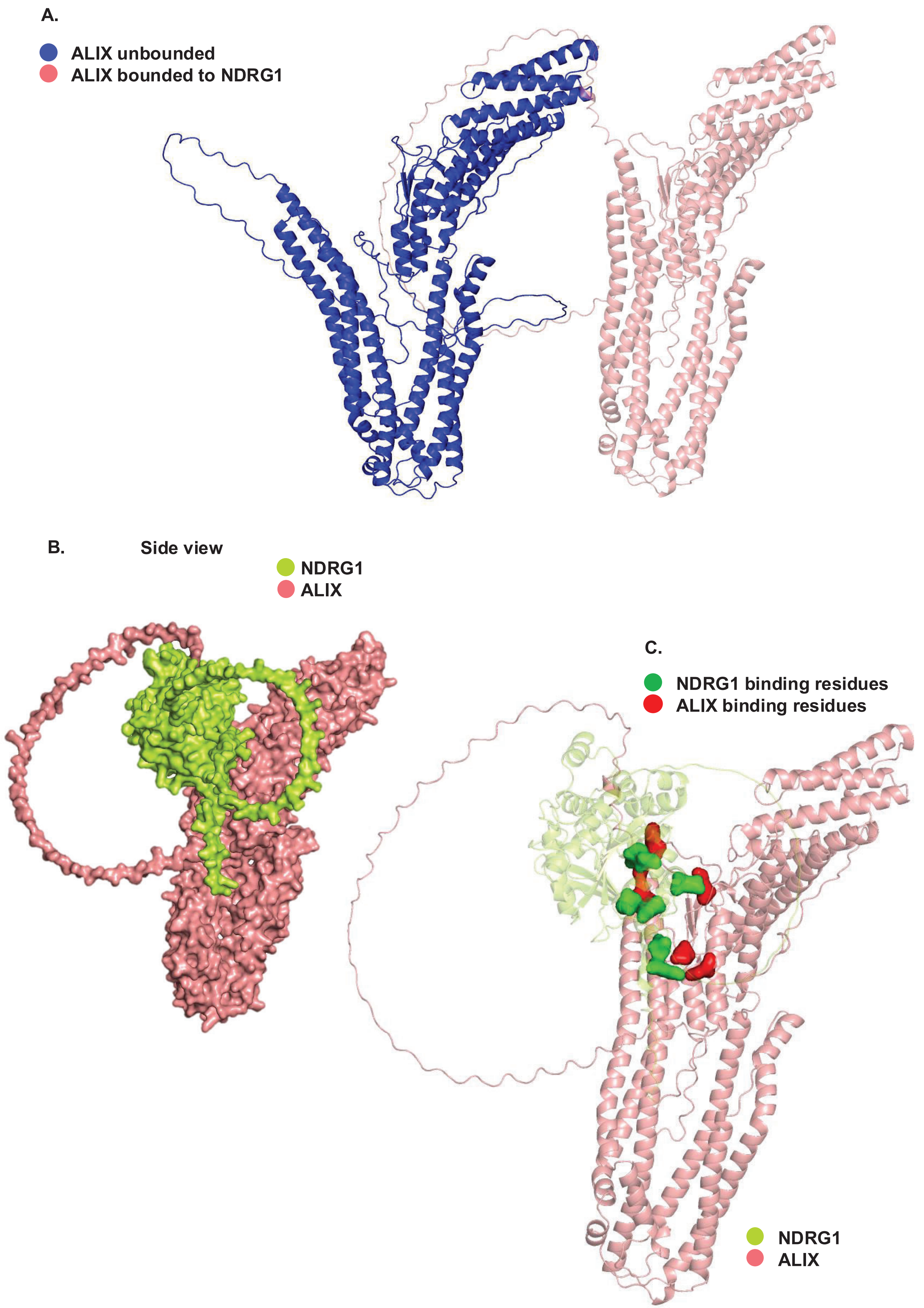


Supplemental Figure 5

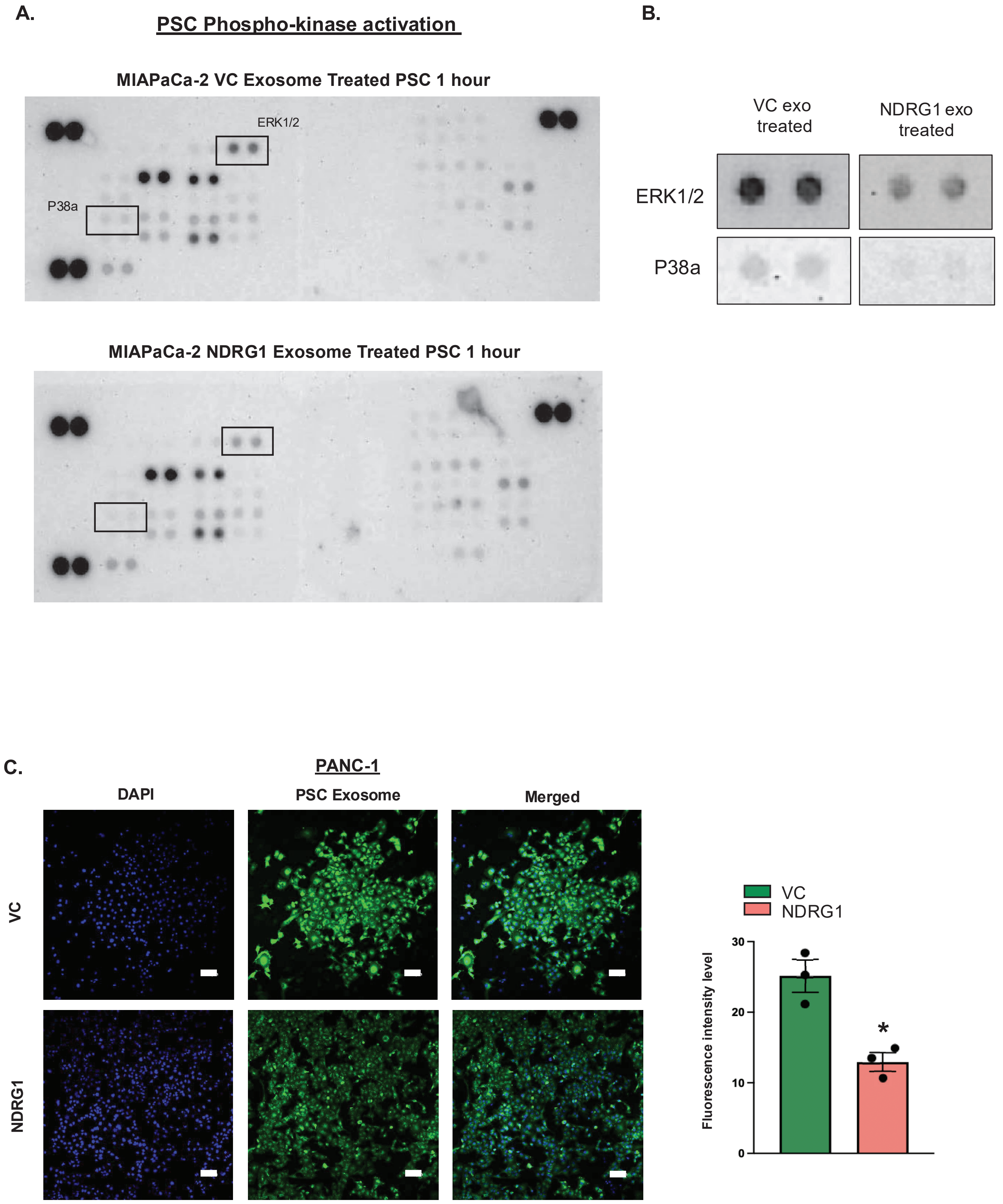


Supplemental Figure 6
